# Narratives: fMRI data for evaluating models of naturalistic language comprehension

**DOI:** 10.1101/2020.12.23.424091

**Authors:** Samuel A. Nastase, Yun-Fei Liu, Hanna Hillman, Asieh Zadbood, Liat Hasenfratz, Neggin Keshavarzian, Janice Chen, Christopher J. Honey, Yaara Yeshurun, Mor Regev, Mai Nguyen, Claire H. C. Chang, Christopher Baldassano, Olga Lositsky, Erez Simony, Michael A. Chow, Yuan Chang Leong, Paula P. Brooks, Emily Micciche, Gina Choe, Ariel Goldstein, Tamara Vanderwal, Yaroslav O. Halchenko, Kenneth A. Norman, Uri Hasson

## Abstract

The “Narratives” collection aggregates a variety of functional MRI datasets collected while human subjects listened to naturalistic spoken stories. The current release includes 345 subjects, 891 functional scans, and 27 diverse stories of varying duration totaling ~4.6 hours of unique stimuli (~43,000 words). This data collection is well-suited for naturalistic neuroimaging analysis, and is intended to serve as a benchmark for models of language and narrative comprehension. We provide standardized MRI data accompanied by rich metadata, preprocessed versions of the data ready for immediate use, and the spoken story stimuli with time-stamped phoneme- and word-level transcripts. All code and data are publicly available with full provenance in keeping with current best practices in transparent and reproducible neuroimaging.

## Background and summary

We use language to build a shared understanding of the world. In speaking, we package certain brain states into a sequence of linguistic elements that can be transmitted verbally, through vibrations in the air; in listening, we expand a verbal transmission into the intended brain states, bringing our own experiences to bear on the interpretation^1^. The neural machinery supporting this language faculty presents a particular challenge for neuroscience. Certain aspects of language are uniquely human^2^, limiting the efficacy of nonhuman animal models and restricting the use of invasive methods (for example, only humans recursively combine linguistic elements into complex, hierarchical expressions with long-range dependencies^3,4^). Language is also very dynamic and contextualized. Linguistic narratives evolve over time: the meaning of any given element depends in part on the history of elements preceding it and certain elements may retroactively resolve the ambiguity of preceding elements. This makes it difficult to decompose natural language using traditional experimental manipulations^5–7^.

Noninvasive neuroimaging tools, such as functional MRI, have laid the groundwork for a neurobiology of language comprehension^8–14^. Functional MRI measures local fluctuations in blood oxygenation—referred to as the blood-oxygen-level-dependent (BOLD) signal—associated with changes in neural activity over the course of an experimental paradigm^15–18^. Despite relatively poor temporal resolution (e.g. one sample every 1–2 seconds), fMRI allows us to map brain-wide responses to linguistic stimuli at a spatial resolution of millimeters. A large body of prior work has revealed a partially left-lateralized network of temporal and frontal cortices encoding acoustic-phonological^19–22^, syntactic^23–25^, and semantic features^26,27^ of linguistic stimuli (with considerable individual variability^28–30^). This work has historically been grounded in highly-controlled experimental paradigms focusing on isolated phonemes^31,32^, single words^33–35^, or contrasting sentence manipulations^36–41^ (with some exceptions^42,43^). While these studies have provided a tremendous number of insights, the vast majority rely on highly-controlled experimental manipulations with limited generalizability to natural language^5,44^. Recent work, however, has begun extending our understanding to more ecological contexts using naturalistic text^45^ and speech^46^.

In parallel, the machine learning community has made tremendous advances in natural language processing^47,48^. Neurally-inspired computational models are beginning to excel at complex linguistic tasks such as word-prediction, summarization, translation, and question-answering^49,50^. Rather than using symbolic lexical representations and syntactic trees, these models typically rely on vectorial representations of linguistic content^51^: linguistic elements that are similar in some respect are encoded nearer to each other in a continuous embedding space, and seemingly complex linguistic relations can be recovered using relatively simple arithmetic operations^52,53^. In contrast to experimental traditions in linguistics and neuroscience, machine learning has embraced complex, high-dimensional models trained on enormous corpora of real-world text, emphasizing predictive power over interpretability^54–56^.

We expect that a reconvergence of these research trajectories supported by “big” neural data will be mutually beneficial^44,57^. Public, well-curated benchmark datasets can both accelerate research and serve as useful didactic tools (e.g. MNIST^58^, CIFAR^59^. Furthermore, there is a societal benefit to sharing human brain data: fMRI data are expensive to acquire and the reuse of publicly shared fMRI data is estimated to have saved billions in public funding^60^. Public data also receive much greater scrutiny, with the potential to reveal (and rectify) “bugs” in the data or metadata. Although public datasets released by large consortia have propelled exploratory research forward^61,62^, relatively few of these include naturalistic stimuli (cf. movie-viewing paradigms in Cam-CAN^63,64^, HBN^65^, HCP S1200^62^). On the other hand, maturing standards and infrastructure^66–68^ have enabled increasingly widespread sharing of smaller datasets from the “long tail” of human neuroimaging research^69^. We are beginning to see a proliferation of public neuroimaging datasets acquired using rich, naturalistic experimental paradigms^45,70–83^. The majority of these datasets are not strictly language-oriented, comprising audiovisual movie stimuli rather than audio-only spoken stories.

With this in mind, we introduce the “Narratives” collection of naturalistic story-listening fMRI data for evaluating models of language^84^. The Narratives collection comprises fMRI data collected over the course of seven years by the Hasson and Norman Labs at the Princeton Neuroscience Institute while participants listened to 27 spoken story stimuli ranging from ~3 minutes to ~56 minutes for a total of ~4.6 hours of unique stimuli (~43,000 words; Table 1). The collection currently includes 345 unique subjects contributing 891 functional scans with accompanying anatomical data and metadata. In addition to the MRI data, we provide demographic data and comprehension scores where available. Finally, we provide the auditory story stimuli, as well as time-stamped phoneme- and word-level transcripts in hopes of accelerating analysis of the linguistic content of the data. The data are standardized according to the Brain Imaging Data Structure^85^ (BIDS 1.2.1; https://bids.neuroimaging.io/; RRID:SCR_016124), and are publicly available via OpenNeuro^86^ (https://openneuro.org/; RRID:SCR_005031): https://openneuro.org/datasets/ds002345. Derivatives of the data, including preprocessed versions of the data and stimulus annotations, are available with transparent provenance via DataLad^87,88^ (https://www.datalad.org/; RRID:SCR_003931): http://datasets.datalad.org/?dir=/labs/hasson/narratives.

**Table 1.**
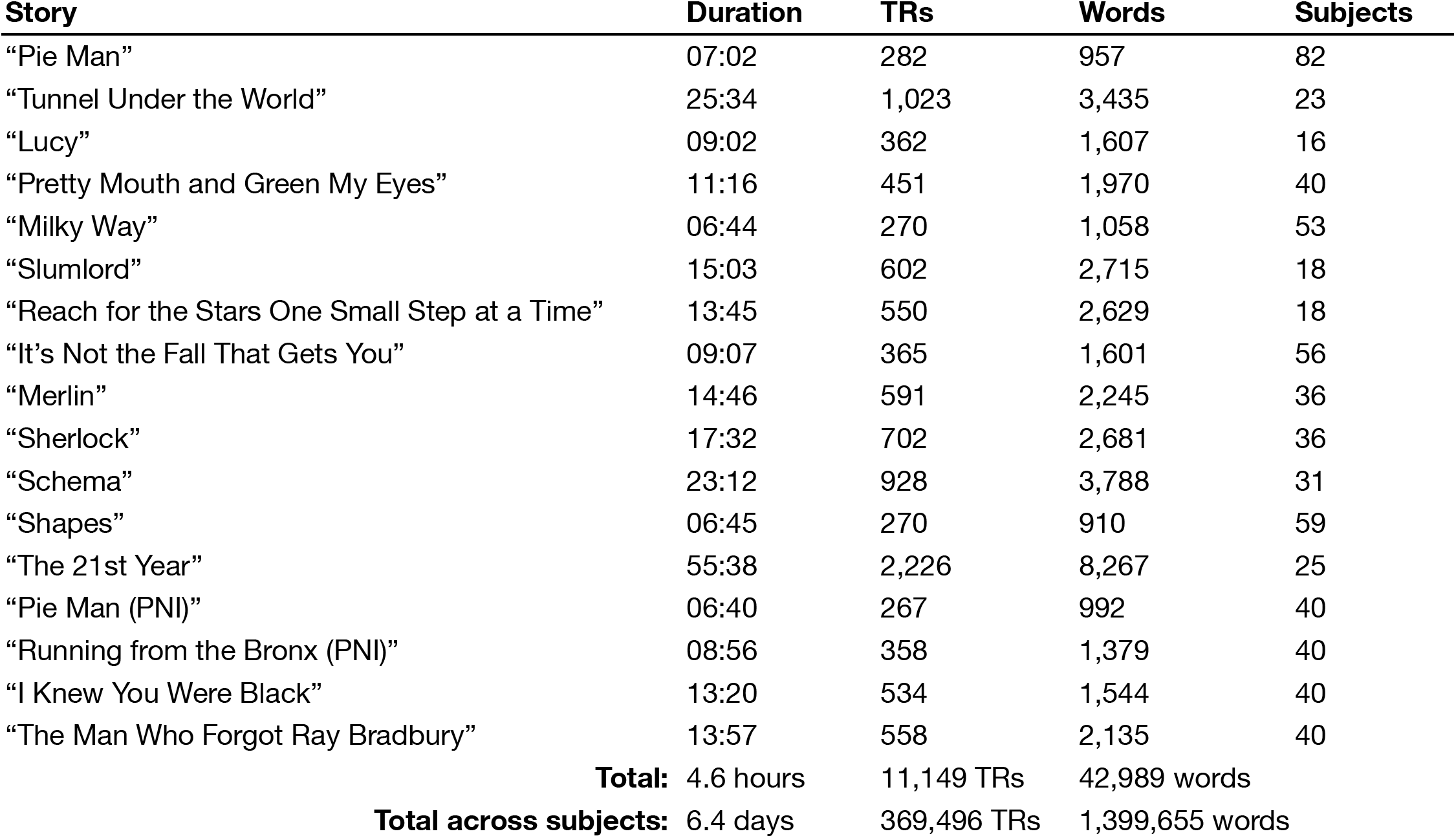
The “Narratives” datasets summarized in terms of duration, word count, and sample size. Some subjects participated in multiple experiments resulting in overlapping samples. Note that for the “Milky Way” and “Shapes” datasets, we tabulate the duration and word count only once for closely related experimental manipulations, respectively (reflected in the total durations at bottom). For the “Schema” dataset, we indicate the sum of the duration and word counts across eight brief stories. We do not include the temporally scrambled versions of the “It’s Not the Fall That Gets You Dataset” in the duration and word totals.

We believe these data have considerable potential for reuse because naturalistic spoken narratives are rich enough to support testing of a wide range of meaningful hypotheses about language and the brain^5,7,44,89–94^. These hypotheses can be formalized as quantitative models and evaluated against the brain data^95,96^ (Fig. 1a). For example, the Narratives data are particularly well-suited for evaluating models capturing linguistic content ranging from lower-level acoustic features^97–99^ to higher-level semantic features^46,100–102^. More broadly, naturalistic data of this sort can be useful for evaluating shared information across subjects^103,104^, individual differences^76,105–107^, algorithms for functional alignment algorithms for functional alignment^80,108–112^, models of event segmentation and narrative context^113–117^, and functional network organization^118–121^. In the following, we describe the Narratives data collection and provide several perspectives on data quality.

**Fig. 1.**
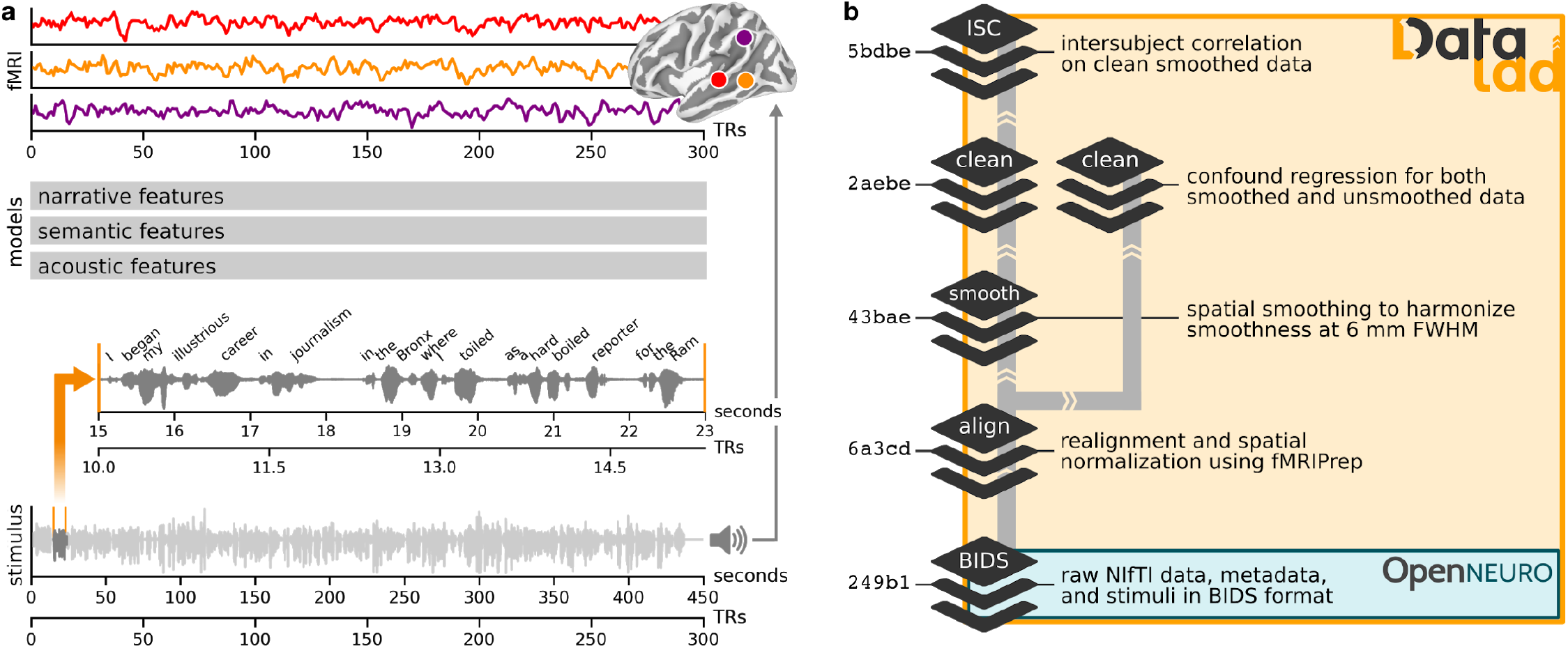
Schematic depiction of the naturalistic story-listening paradigm and data provenance. (**a**) At bottom, the full auditory story stimulus “Pie Man” by Jim O’Grady is plotted as a waveform of varying amplitude (*y*-axis) over the duration of 450 seconds (*x*-axis) corresponding to 300 fMRI volumes sampled at a TR of 1.5 seconds. An example clip (marked by vertical orange lines) is expanded and accompanied by the time-stamped word annotation (“I began my illustrious career in journalism in the Bronx, where I worked as a hard-boiled reporter for the Ram…”). The story stimulus can be described according to a variety of models; for example, acoustic, semantic, or narrative features can be extracted from or assigned to the stimulus. In a prediction or model-comparison framework, these models serve as formal hypotheses linking the stimulus to brain responses. At top, preprocessed fMRI response time-series from three example voxels for an example subject are plotted for the full duration of the story stimulus (*x*-axis: fMRI signal magnitude; *y*-axis: scan duration in TRs; red: early auditory cortex; orange: auditory association cortex; purple: temporoparietal junction). See the `plot_stim.py` script in the `code/` directory for details. (**b**) At bottom, MRI data, metadata, and stimuli are formatted according to the BIDS standard and publicly available via OpenNeuro. All derivative data are version-controlled and publicly available via DataLad. The schematic preprocessing workflow includes the following steps: realignment, susceptibility distortion correction, and spatial normalization with fMRIPrep; unsmoothed and spatially smoothed workflows proceed in parallel; confound regression to mitigate artifacts from head motion and physiological noise; as well as intersubject correlation (ISC) analyses used for quality control in this manuscript. Each stage of the processing workflow is publicly available and indexed by a commit hash (left) providing a full, interactive history of the data collection. This schematic is intended to provide a high-level summary and does not capture the full provenance in detail; for example, derivatives from MRIQC are also included in the public release alongside other derivatives (but are not depicted here).

## Methods

### Participants

Data were collected over the course of roughly seven years, from October, 2011 to September, 2018. Participants were recruited from the Princeton University student body as well as non-university-affiliated members of the broader community in Princeton, NJ. All participants provided informed, written consent prior to data collection in accordance with experimental procedures approved by Princeton University Institutional Review Board. Across all datasets, 345 adults participated in data collection (ages 18–53 years, mean age 22.2 ± 5.1 years, 204 reported female). Demographics for each dataset are reported in the “Narrative datasets” section. Both native and non-native English speakers are included in the dataset, but all subjects reported fluency in English (our records do not contain detailed information on fluency in other languages). All subjects reported having normal hearing and no history of neurological disorders.

### Experimental procedures

Upon arriving at the fMRI facility, participants first completed a simple demographics questionnaire, as well as a comprehensive MRI screening form. Participants were instructed to listen and pay attention to the story stimulus, remain still, and keep their eyes open. In some cases, subject wakefulness was monitored in real-time using an eye-tracker. Stimulus presentation was implemented using either Psychtoolbox^122,123^ or PsychoPy^124–126^. In some cases a centrally-located fixation cross or dot was presented throughout the story stimulus; however, participants were not instructed to maintain fixation. Auditory story stimuli were delivered via MRI-compatible insert earphones (Sensimetrics, Model S14); headphones or foam padding were placed over the earphones to reduce scanner noise. In most cases, a volume check was performed to ensure that subjects could comfortably hear the auditory stimulus over the MRI acquisition noise prior to data collection. A sample audio stimulus was played during an EPI sequence (that was later discarded) and the volume was adjusted by either the experimenter or subject until the subject reported being able to comfortably hear and understand the stimulus. In the `staging/` directory on GitHub, we provide an example stimulus presentation PsychoPy script used with the “Pie Man (PNI)”, “Running from the Bronx (PNI)”, “I Knew You Were Black”, and “The Man Who Forgot Ray Bradbury” stories (`story_presentation.py`), as well as an example volume-check script (`soundcheck_presentation.py`; see “Code availability”).

For many datasets in this collection, story scans were accompanied by additional functional scans including different stories or other experimental paradigms, as well as auxiliary anatomical scans. In all cases, an entire story stimulus was presented in a single scanning run; however, in some cases multiple independent stories were collected in a single scanning run (the “Slumlord” and “Reach for the Stars One Small Step at a Time” stories, and the “Schema” stories; see the “Narrative datasets” section below). In scanning sessions with multiple stories and scanning runs, participants were free to request volume adjustments between scans (although this was uncommon). Participants were debriefed as to the purpose of a given experiment at the end of a scanning session, and in some cases filled out follow-up questionnaires evaluating their comprehension of the stimuli. Specifical procedural details for each dataset are described in the “Narrative datasets” section below.

### Stimuli

To add to the scientific value of the neuroimaging data, we provide waveform (WAV) audio files containing the spoken story stimulus for each dataset (e.g. `pieman_audio.wav`). Audio files were edited using the open source Audacity software (https://www.audacityteam.org). These audio files served as stimuli for the publicly-funded psychological and neuroscientific studies described herein. The stimuli are intended for non-profit, non-commercial scholarly use—principally feature extraction—under “fair use” or “fair dealing” provisions; (re)sharing and (re)use of these media files is intended to respect these guidelines^127^. Story stimuli span a variety of media, including commercially-produced radio and internet broadcasts, authors and actors reading written works, professional storytellers performing in front of live audiences, and subjects verbally recalling previous events (in the scanner). Manually written transcripts are provided with each story stimulus. The specific story stimuli for each dataset are described in more detail in the “Narrative datasets” section below. In total, the auditory stories sum to roughly 4.6 hours of unique stimuli corresponding to 11,149 TRs (excluding TRs acquired with no auditory story stimulus). Concatenating across all subjects, this amounts to roughly 6.4 days worth of story-listening fMRI data, or 369,496 TRs.

By including all stimuli in this data release, researchers can extract their choice of linguistic features for model evaluation^7,128^, opening the door to many novel scientific questions. To kick-start this process, we used Gentle 0.10.1^129^ to create time-stamped phoneme- and word-level transcripts for each story stimulus in an automated fashion. Gentle is a robust, lenient forced-alignment algorithm that relies on the free and open source Kaldi automated speech recognition software^130^ and the Fisher English Corpus^131^. The initial (non-time-stamped) written transcripts were manually transcribed by the authors and supplied to the Gentle forced-alignment algorithm alongside the audio file. The first-pass output of the forced-alignment algorithm was visually inspected and any errors in the original transcripts were manually corrected; then the corrected transcripts were resubmitted to the forced-alignment algorithm to generate the final time-stamped transcripts.

The Gentle forced-alignment software generates two principal outputs. First, a simple word-level transcript with onset and offset timing for each word is saved as a tabular CSV file. Each row in the CSV file corresponds to a word. The CSV file contains four columns (no header) where the first column indicates the word from the written transcript, the second column indicates the corresponding word in the vocabulary, the third column indicates the word onset timing from the beginning of the stimulus file, and the fourth column indicates the word offset timing. Second, Gentle generates a more detailed JSON file containing the full written transcript followed by a list of word entries where each entry contains fields indicating the word (“word”) and whether word was found in the vocabulary (“alignedWord”), the onset (“start”) and offset (“end”) timing of the word referenced to the beginning of the audio file, the duration of each phone comprising that word (“phones” containing “phone” and “duration”), and a field (“case”) indicating whether the word was correctly localized in the audio signal. Words that were not found in the vocabulary are indicated by “<unk>” for “unknown” in the second column of the CSV file or in the “alignedWord” field of the JSON file. Words that are not successfully localized in the audio file receive their own row but contain no timing information in the CSV file, and are marked as “not-found-in-audio” in the “case” field of the JSON file. All timing information generated by Gentle is indexed to the beginning of the stimulus audio file. This should be referenced against the BIDS-formatted `events.tsv` files accompanying each scan describing when the story stimulus began relative to the beginning of the scan (this varies across datasets as described in the “Narratives datasets” section below). Some word counts may be slightly inflated by the inclusion of disfluencies (e.g. “uh”) in the transcripts. Transcripts for the story stimuli contain 42,989 words in total across all stories; 789 words (1.8%) were not successfully localized by the forced-alignment algorithm and 651 words (1.5%) were not found in the vocabulary (see the `get_words.py` script in the `code/` directory). Concatenating across all subjects yields 1,399,655 words occurring over the course of 369,496 TRs.

Gentle packages the audio file, written transcript, CSV and JSON files with an HTML file for viewing in a browser that allows for interactively listening to the audio file while each word is highlighted (with its corresponding phones) when it occurs in the audio. Note that our forced-alignment procedure using Gentle was fully automated and did not include any manual intervention. That said, the time-stamped transcripts are not perfect—speech disfluencies, multiple voices speaking simultaneously, sound effects, and scanner noise provide a challenge for automated transcription. We hope this annotation can serve as a starting point, and expect that better transcriptions will be created in the future. We invite researchers who derive additional features and annotations from the stimuli to contact the corresponding author and we will help incorporate these annotations into the publicly available dataset.

### MRI data acquisition

All datasets were collected at the Princeton Neuroscience Institute Scully Center for Neuroimaging. Acquisition parameters are reproduced below in compliance with the guidelines put forward by the Organization for Human Brain Mapping (OHBM) Committee on Best Practices in Data Analysis and Sharing^132^ (COBIDAS), and are also included in the BIDS-formatted JSON metadata files accompanying each scan. All studies used a repetition time (TR) of 1.5 seconds. Several groups of datasets differ in acquisition parameters due to both experimental specifications and the span of years over which datasets were acquired; for example, several of the newer datasets were acquired on a newer scanner and use multiband acceleration to achieve greater spatial resolution (e.g. for multivariate pattern analysis) while maintaining the same temporal resolution. The “Pie Man”, “Tunnel Under the World”, “Lucy”, “Pretty Mouth and Green My Eyes”, “Milky Way”, “Slumlord”, “Reach for the Stars One Small Step at a Time”, “It’s Not the Fall that Gets You”, “Merlin”, “Sherlock”, and “The 21st Year” datasets were collected on a 3T Siemens Magnetom Skyra (Erlangen, Germany) with a 20-channel phased-array head coil using the following acquisition parameters. Functional BOLD images were acquired in an interleaved fashion using gradient-echo echo-planar imaging (EPI) with an in-plane acceleration factor of 2 using GRAPPA: TR/TE = 1500/28 ms, flip angle = 64°, bandwidth = 1445 Hz/Px, in-plane resolution = 3 × 3 mm, slice thickness = 4 mm, matrix size = 64 × 64, FoV = 192 × 192 mm, 27 axial slices with roughly full brain coverage and no gap, anterior–posterior phase encoding, prescan normalization, fat suppression. In cases where full brain coverage was not attainable, inferior extremities were typically excluded (e.g. cerebellum, brainstem) to maximize coverage of the cerebral cortex. At the beginning of each run, three dummy scans were acquired and discarded by the scanner to allow for signal stabilization. T1-weighted structural images were acquired using a high-resolution single-shot MPRAGE sequence with an in-plane acceleration factor of 2 using GRAPPA: TR/TE/TI = 2300/3.08/900 ms, flip angle = 9°, bandwidth = 240 Hz/Px, in-plane resolution 0.859 × 0.859 mm, slice thickness 0.9 mm, matrix size = 256 × 256, FoV = 172.8 × 220 × 220 mm, 192 sagittal slices, ascending acquisition, anterior–posterior phase encoding, prescan normalization, no fat suppression, 7 minutes 21 seconds total acquisition time.

The “Schema” and “Shapes” datasets were collected on a 3T Siemens Magnetom Prisma with a 64-channel head coil using the following acquisition parameters. Functional images were acquired in an interleaved fashion using gradient-echo EPI with a multiband (simultaneous multi-slice; SMS) acceleration factor of 4 using blipped CAIPIRINHA and no in-plane acceleration: TR/TE 1500/39 ms, flip angle = 50°, bandwidth = 1240 Hz/Px, in-plane resolution = 2.0 × 2.0 mm, slice thickness 2.0 mm, matrix size = 96 × 96, FoV = 192 × 192 mm, 60 axial slices with full brain coverage and no gap, anterior–posterior phase encoding, 6/8 partial Fourier, no prescan normalization, fat suppression, three dummy scans. T1-weighted structural images were acquired using a high-resolution single-shot MPRAGE sequence with an in-plane acceleration factor of 2 using GRAPPA: TR/TE/TI = 2530/2.67/1200 ms, flip angle = 7°, bandwidth = 200 Hz/Px, in-plane resolution 1.0 × 1.0 mm, slice thickness 1.0 mm, matrix size = 256 × 256, FoV = 176 × 256 × 256 mm, 176 sagittal slices, ascending acquisition, no fat suppression, 5 minutes 52 seconds total acquisition time.

The “Pie Man (PNI),” “Running from the Bronx,” “I Knew You Were Black,” and “The Man Who Forgot Ray Bradbury” datasets were collected on the same 3T Siemens Magnetom Prisma with a 64-channel head coil using different acquisition parameters. Functional images were acquired in an interleaved fashion using gradient-echo EPI with a multiband acceleration factor of 3 using blipped CAIPIRINHA and no in-plane acceleration: TR/TE 1500/31 ms, flip angle = 67°, bandwidth = 2480 Hz/Px, in-plane resolution = 2.5 × 2.5 mm, slice thickness 2.5 mm, matrix size = 96 × 96, FoV = 240 × 240 mm, 48 axial slices with full brain coverage and no gap, anterior–posterior phase encoding, prescan normalization, fat suppression, three dummy scans. T1-weighted structural images were acquired using a high-resolution single-shot MPRAGE sequence with an in-plane acceleration factor of 2 using GRAPPA: TR/TE/TI = 2530/3.3/1100 ms, flip angle = 7°, bandwidth = 200 Hz/Px, in-plane resolution 1.0 × 1.0 mm, slice thickness 1.0 mm, matrix size = 256 × 256, FoV = 176 × 256 × 256 mm, 176 sagittal slices, ascending acquisition, no fat suppression, prescan normalization, 5 minutes 53 seconds total acquisition time. T2-weighted structural images were acquired using a high-resolution single-shot MPRAGE sequence with an in-plane acceleration factor of 2 using GRAPPA: TR/TE = 3200/428 ms, flip angle = 120°, bandwidth = 200 Hz/Px, in-plane resolution 1.0 × 1.0 mm, slice thickness 1.0 mm, matrix size = 256 × 256, FoV = 176 × 256 × 256 mm, 176 sagittal slices, interleaved acquisition, no prescan normalization, no fat suppression, 4 minutes 40 seconds total acquisition time.

### MRI preprocessing

Anatomical images were de-faced using the automated de-facing software pydeface 2.0.0^133^ prior to further processing (using the `run_pydeface.py` script in the `code/` directory). MRI data were subsequently preprocessed using fMRIPrep 20.0.5^134,135^ (RRID:SCR_016216; using the `run_fmriprep.sh` script in the `code/` directory). FMRIPrep is a containerized, automated tool based on Nipype 1.4.2^136,137^ (RRID:SCR_002502) that adaptively adjusts to idiosyncrasies of the dataset (as captured by the metadata) to apply the best-in-breed preprocessing workflow. Many internal operations of fMRIPrep functional processing workflow use Nilearn 0.6.2^138^ (RRID:SCR_001362). For more details of the pipeline, see the section corresponding to workflows in fMRIPrep’s documentation. The containerized fMRIPrep software was deployed using Singularity 3.5.2-1.1.sdl7^139^. The fMRIPrep Singularity image can be built from Docker Hub (https://hub.docker.com/r/poldracklab/fmriprep/; e.g. `singularity build fmriprep-20.0.5.simg docker://poldracklab/fmriprep:20.0.5`). The fMRIPrep outputs and visualization can be found in the `fmriprep/` directory in `derivatives/` available via the DataLad release. The fMRIPrep workflow produces two principal outputs: (*a*) the functional time series data in one more output space (e.g. MNI space), and (*b*) a collection of confound variables for each functional scan. In the following, we describe fMRIPrep’s anatomical and functional workflows, as well as subsequent spatial smoothing and confound regression implemented in AFNI 19.3.0^140,141^ (RRID:SCR_005927).

The anatomical MRI T1-weighted (T1w) images were corrected for intensity non-uniformity with `N4BiasFieldCorrection`^142^, distributed with ANTs 2.2.0^143^ (RRID:SCR_004757), and used as T1w-reference throughout the workflow. The T1w-reference was then skull-stripped with a Nipype implementation of the `antsBrainExtraction.sh` (from ANTs) using the OASIS30ANTs as the target template. Brain tissue segmentation of cerebrospinal fluid (CSF), white-matter (WM), and gray-matter (GM) was performed on the brain-extracted T1w using `fast`^144^ (FSL 5.0.9; RRID:SCR_002823). Brain surfaces were reconstructed using `recon-all`^145,146^ (FreeSurfer 6.0.1; RRID:SCR_001847), and the brain mask estimated previously was refined with a custom variation of the method to reconcile ANTs-derived and FreeSurfer-derived segmentations of the cortical gray-matter from Mindboggle^147^ (RRID:SCR_002438). Volume-based spatial normalization to two commonly-used standard spaces (*MNI152NLin2009cAsym, MNI152NLin6Asym*) was performed through nonlinear registration with `antsRegistration` (ANTs 2.2.0) using brain-extracted versions of both T1w reference and the T1w template. The following two volumetric templates were selected for spatial normalization and deployed using TemplateFlow^148^: (*a*) *ICBM 152 Nonlinear Asymmetrical Template Version 2009c*^149^ (RRID:SCR_008796; TemplateFlow ID: MNI152NLin2009cAsym), and (*b*) *FSL’s MNI ICBM 152 Non-linear 6th Generation Asymmetric Average Brain Stereotaxic Registration Model*^150^ (RRID:SCR_002823; TemplateFlow ID: MNI152NLin6Asym). Surface-based normalization based on nonlinear registration of sulcal curvature was applied using the following three surface templates^151^ (FreeSurfer reconstruction nomenclature): *fsaverage, fsaverage6, fsaverage5*.

The functional MRI data were preprocessed in the following way. First, a reference volume and its skull-stripped version were generated using a custom methodology of fMRIPrep. A deformation field to correct for susceptibility distortions was estimated using fMRIPrep’s fieldmap-less approach. The deformation field results from co-registering the BOLD reference to the same-subject T1w-reference with its intensity inverted^152,153^. Registration was performed with `antsRegistration` (ANTs 2.2.0), and the process was regularized by constraining deformation to be nonzero only along the phase-encoding direction, and modulated with an average fieldmap template^154^. Based on the estimated susceptibility distortion, a corrected EPI reference was calculated for more accurate co-registration with the anatomical reference. The BOLD reference was then co-registered to the T1w reference using `bbregister` (FreeSurfer 6.0.1), which implements boundary-based registration^155^. Co-registration was configured with six degrees of freedom. Head-motion parameters with respect to the BOLD reference (transformation matrices, and six corresponding rotation and translation parameters) are estimated before any spatiotemporal filtering using `mcflirt` (FSL 5.0.9)^156–158^. BOLD runs were slice-time corrected using `3dTshift` from AFNI 20160207^159^. The BOLD time-series were resampled onto the following surfaces: *fsaverage, fsaverage6, fsaverage5*. The BOLD time-series (including slice-timing correction when applied) were resampled onto their original, native space by applying a single, composite transform to correct for head-motion and susceptibility distortions. These resampled BOLD time-series are referred to as preprocessed BOLD in original space, or just preprocessed BOLD. The BOLD time-series were resampled into two volumetric standard spaces, correspondingly generating the following spatially-normalized, preprocessed BOLD runs: *MNI152NLin2009cAsym, MNI152NLin6Asym*. A reference volume and its skull-stripped version were first generated using a custom methodology of fMRIPrep. All resamplings were performed with a single interpolation step by composing all the pertinent transformations (i.e. head-motion transform matrices, susceptibility distortion correction, and co-registrations to anatomical and output spaces). Gridded (volumetric) resamplings were performed using `antsApplyTransforms` (ANTs 2.2.0), configured with Lanczos interpolation to minimize the smoothing effects of other kernels^160^. Non-gridded (surface) resamplings were performed using `mri_vol2surf` (FreeSurfer 6.0.1).

Several confounding time-series were calculated based on the preprocessed BOLD: framewise displacement (FD), DVARS, and three region-wise global signals. FD and DVARS are calculated for each functional run, both using their implementations in Nipype^161^. The three global signals are extracted within the CSF, the WM, and the whole-brain masks. Additionally, a set of physiological regressors were extracted to allow for component-based noise correction (CompCor)^162^. Principal components are estimated after high-pass filtering the preprocessed BOLD time-series (using a discrete cosine filter with 128 s cut-off) for the two CompCor variants: temporal (tCompCor) and anatomical (aCompCor). The tCompCor components are then calculated from the top 5% variable voxels within a mask covering the subcortical regions. This subcortical mask is obtained by heavily eroding the brain mask, which ensures it does not include cortical GM regions. For aCompCor, components are calculated within the intersection of the aforementioned mask and the union of CSF and WM masks calculated in T1w space, after their projection to the native space of each functional run (using the inverse BOLD-to-T1w transformation). Components are also calculated separately within the WM and CSF masks. For each CompCor decomposition, the *k* components with the largest singular values are retained, such that the retained components’ time series are sufficient to explain 50 percent of variance across the nuisance mask (CSF, WM, combined, or temporal). The remaining components are dropped from consideration. The head-motion estimates calculated in the correction step were also placed within the corresponding confounds file. The confound time series derived from head motion estimates and global signals were expanded with the inclusion of temporal derivatives and quadratic terms for each^163^. Frames that exceeded a threshold of 0.5 mm FD or 1.5 standardised DVARS were annotated as motion outliers. All of these confound variables are provided with the dataset for researchers to use as they see fit. HTML files with quality control visualizations output by fMRIPrep are available via DataLad.

We provide spatially smoothed and non-smoothed versions of the preprocessed functional data returned by fMRIPrep (smoothing was implemented using the `run_smoothing.py` script in the `code/` directory). Analyses requiring voxelwise correspondence across subjects, such as ISC analysis^164^, can benefit from spatial smoothing due to variability in functional–anatomical correspondence across individuals—at the expense of spatial specificity (finer-grained intersubject functional correspondence can be achieved using hyperalignment rather than spatial smoothing^80,112,165^). The smoothed and non-smoothed outputs (and subsequent analyses) can be found in the `afni-smooth/` and `afni-nosmooth/` directories in `derivatives/` available via DataLad. To smooth the volumetric functional images, we used `3dBlurToFWHM` in AFNI 19.3.0^140,141^ which iteratively measures the global smoothness (ratio of variance of first differences across voxels to global variance) and local smoothness (estimated within 4 mm spheres), then applies local smoothing to less-smooth areas until the desired global smoothness is achieved. All smoothing operations were performed within a brain mask to ensure that non-brain values were not smoothed into the functional data (see the `brain_masks.py` script in the `code/` directory). For surface-based functional data, we applied smoothing `SurfSmooth` in AFNI, which uses a similar iterative algorithm for smoothing surface data according to geodesic distance on the cortical mantle^166,167^. In both cases, data were smoothed until a target global smoothness of 6 mm FWHM was met (i.e. 2–3 times original voxel sizes^168^). Equalizing smoothness is critical to harmonize data acquired across different scanners and protocols^169^.

We next temporally filtered the functional data to mitigate the effects of confounding variables. Unlike traditional task fMRI experiments with a well-defined event structure, the goal of regression was not to estimate regression coefficients for any given experimental conditions; rather, similar to resting-state functional connectivity analysis, the goal of regression was to model nuisance variables, resulting in a “clean” residual time series. However, unlike conventional resting-state paradigms, naturalistic stimuli enable intersubject analyses, which are less sensitive to idiosyncratic noises than within-subject functional connectivity analysis typically used with resting-state data^118,170^. With this in mind, we used a modest confound regression model informed by the rich literature on confound regression for resting-state functional connectivity^171,172^. AFNI’s `3dTproject` was used to regress out the following nuisance variables (via the `extract_confounds.py` and `run_regression.py` scripts in the `code/` directory): six head motion parameters (three translation, three rotation), the first five principal component time series from an eroded CSF and a white matter mask^162,173^, cosine bases for high-pass filtering (using a discrete cosine filter with cutoff: 128 s, or ~.0078 Hz), and first- and second-order detrending polynomials. These variables were included in a single regression model to avoid reintroducing artifacts by sequential filtering^174^. The scripts used to perform this regression and the residual time series are provided with this data release. This processing workflow ultimately yields smoothed and non-smoothed versions of the “clean” functional time series data in several volumetric and surface-based standard spaces.

### Computing environment

In addition to software mentioned elsewhere in this manuscript, all data processing relied heavily on the free, open source GNU/Linux ecosystem and NeuroDebian distribution^175,176^ (https://neuro.debian.net/; RRID:SCR_004401), as well as the Anaconda distribution (https://www.anaconda.com/) and conda package manager (https://docs.conda.io/en/latest/; RRID:SCR_018317). Many analyses relied on scientific computing software in Python (https://www.python.org/; RRID:SCR_008394), including NumPy^177,178^ (http://www.numpy.org/; RRID:SCR_008633), SciPy^179,180^ (https://www.scipy.org/; RRID:SCR_008058), Pandas^181^ (https://pandas.pydata.org/; RRID:SCR_018214), NiBabel^182^ (https://nipy.org/nibabel/; RRID:SCR_002498), IPython^183^ (http://ipython.org/; RRID:SCR_001658), and Jupyter^184^ (https://jupyter.org/; RRID:SCR_018315), as well as Singularity containerization^139^ (https://sylabs.io/docs/) and the Slurm workload manager^185^ (https://slurm.schedmd.com/). All surface-based MRI data were visualized using SUMA^186,187^ (RRID:SCR_005927). All other figures were created using Matplotlib^188^ (http://matplotlib.org/; RRID:SCR_008624), seaborn (http://seaborn.pydata.org/; RRID:SCR_018132), GIMP (http://www.gimp.org/; RRID:SCR_003182), and Inkscape (https://inkscape.org/; RRID:SCR_014479). The “Narratives” data were processed on a Springdale Linux 7.9 (Verona) system based on the Red Hat Enterprise Linux distribution (https://springdale.math.ias.edu/). An `environment.yml` file specifying the conda environment used to process the data is included in the `staging/` directory on GitHub (as well as a more flexible cross-platform `environment-flexible.yml` file; see “Code availability”).

### Narrative datasets

Each dataset in the “Narratives” collection is described below. The datasets are listed in roughly chronological order of acquisition. For each dataset, we provide the dataset-specific subject demographics, a summary of the stimulus and timing, as well as details of the experimental procedure or design.

### “Pie Man”

The “Pie Man” dataset was collected between October, 2011 and March, 2013, and comprised 82 participants (ages 18–45 years, mean age 22.5 ± 4.3 years, 45 reported female). The “Pie Man” story was told by Jim O’Grady and recorded live at *The Moth*, a non-profit storytelling event, in New York City in 2008 (freely available at https://themoth.org/stories/pie-man). The “Pie Man” audio stimulus was 450 seconds (7.5 minutes) long and began with 13 seconds of neutral introductory music followed by 2 seconds of silence, such that the story itself started at 0:15 and ended at 7:17, for a duration 422 s, with 13 seconds of silence at the end of the scan. The stimulus was started simultaneously with the acquisition of the first functional MRI volume, and the scans comprise 300 TRs, matching the duration of the stimulus. The transcript for the “Pie Man” story stimulus contains 957 words; 3 words (0.3%) were not successfully localized by the forced-alignment algorithm, and 19 words (2.0%) were not found in the vocabulary (e.g. proper names). The “Pie Man” data were in some cases collected in conjunction with temporally scrambled versions of the “Pie Man” stimulus, as well the “Tunnel Under the World” and “Lucy”datasets among others, and share subjects with these datasets (as specified in the `participants.tsv file`). The current release only includes data collected for the intact story stimulus (rather than the temporally scrambled versions of the stimulus). The following subjects received the “Pie Man” stimulus on two separate occasions (specified by `run-1` or `run-2` in the BIDS file naming convention): `sub-001`, `sub-002`, `sub-003`, `sub-004`, `sub-005`, `sub-006`, `sub-008`, `sub-010`, `sub-011`, `sub-012`, `sub-013`, `sub-014`, `sub-015`, `sub-016`. We recommend excluding subjects `sub-001` (both `run-1` and `run-2`), `sub-013` (`run-2`), `sub-014` (`run-2`), `sub-021`, `sub-022`, `sub-038`, `sub-056`, `sub-068`, and `sub-069` (as specified in the `scan_exclude.json` file in the `code/` directory) based on criteria explained in the “Intersubject correlation” section of “Data validation.” Subsets of the “Pie Man” data have been previously reported in numerous publications^108,114,118,189–197^, and additional datasets not shared here have been collected using the “Pie Man” auditory story stimulus^198–201^. In the filename convention (and figures), “Pie Man” is labeled using the task alias `pieman`.

### “Tunnel Under the World”

The “Tunnel Under the World” dataset was collected between May, 2012, and February, 2013, and comprised 23 participants (ages 18–31 years, mean age 22.5 ± 3.8 years, 14 reported female). The “Tunnel Under the World” science-fiction story was authored by Frederik Pohl in 1955 which was broadcast in 1956 as part of the *X Minus One* series, a collaboration between the National Broadcasting Company and Galaxy Science Fiction magazine (freely available at https://www.oldtimeradiodownloads.com). The “Tunnel Under the World” audio stimulus is 1534 seconds (~25.5 minutes) long. The stimulus was started after the first two functional MRI volumes (2 TRs, 3 seconds) were collected, with ~23 seconds of silence after the stimulus. The functional scans comprise 1040 TRs (1560 seconds), except for subjects `sub-004` and `sub-013` which have 1035 and 1045 TRs respectively. The transcript for “Tunnel Under the World” contains 3,435 words; 126 words (3.7%) were not successfully localized by the forced-alignment algorithm and 88 words (2.6%) were not found in the vocabulary. The “Tunnel Under the World” and “Lucy” datasets contain largely overlapping samples of subjects, though were collected in separate sessions, and were, in many cases, collected alongside “Pie Man” scans (as specified in the `participants.tsv file`). We recommend excluding the “Tunnel Under the World” scans for subjects `sub-004` and `sub-013` (as specified in the `scan_exclude.json` file in the `code/` directory). The “Tunnel Under the World” data have been previously reported by Lositsky and colleagues^202^. In the filename convention, “Tunnel Under the World” is labeled using the task alias `tunnel`.

### “Lucy”

The “Lucy” dataset was collected between October, 2012 and January, 2013 and comprised 16 participants (ages 18–31 years, mean age 22.6 ± 3.9 years, 10 reported female). The “Lucy” story was broadcast by the non-profit WNYC public radio in 2010 (freely available at https://www.wnycstudios.org/story/91705-lucy). The “Lucy” audio stimulus is 542 seconds (~9 minutes) long and was started after the first two functional MRI volumes (2 TRs, 3 seconds) were collected. The functional scans comprise 370 TRs (555 seconds). The transcript for “Lucy” contains 1,607 words; 22 words (1.4%) were not successfully localized by the forced-alignment algorithm and 27 words (1.7%) were not found in the vocabulary. The “Lucy” and “Tunnel Under the World” datasets contain largely overlapping samples of subjects, and were acquired contemporaneously with “Pie Man” data. We recommend excluding the “Lucy” scans for subjects `sub-053` and `sub-065` (as specified in the `scan_exclude.json` file in the `code/` directory). The “Lucy” data have not previously been reported. In the filename convention, “Lucy” is labeled using the task alias `lucy`.

### “Pretty Mouth and Green My Eyes”

The “Pretty Mouth and Green My Eyes” dataset was collected between March, 2013, and October, 2013, and comprised 40 participants (ages 18–34 years, mean age 21.4 ± 3.5 years, 19 reported female). The “Pretty Mouth and Green My Eyes” story was authored by J. D. Salinger for The New Yorker magazine (1951) and subsequently published in the Nine Stories collection (1953). The spoken story stimulus used for data collection was based on an adapted version of the original text that was shorter and included a few additional sentences, and was read by a professional actor. The “Pretty Mouth and Green My Eyes” audio stimulus is 712 seconds (~11.9 minutes) long and began with 18 seconds of neutral introductory music followed by 3 seconds of silence, such that the story started at 0:21 (after 14 TRs) and ended at 11:37, for a duration of 676 seconds (~451 TRs), with 15 seconds (10 TRs) of silence at the end of the scan. The functional scans comprised 475 TRs (712.5 seconds). The transcript for “Pretty Mouth and Green My Eyes” contains 1,970 words; 7 words (0.4%) were not successfully localized by the forced-alignment algorithm and 38 words (1.9%) were not found in the vocabulary.

The “Pretty Mouth and Green My Eyes” stimulus was presented to two groups of subjects in two different narrative contexts according between-subject experimental design: (*a*) in the “affair” group, subjects read a short passage implying that the main character was having an affair; (*b*) in the “paranoia” group, subjects read a short passage implying that the main character’s friend was unjustly paranoid (see Yeshurun et al., 2017, for the full prompts). The two experimental groups were randomly assigned such that there were 20 subjects in each group and each subject only received the stimulus under a single contextual manipulation. The group assignments are indicated in the `participants.tsv file`, and the `scans.tsv` file for each subject. Both groups received the identical story stimulus despite receiving differing contextual prompts. Immediately following the scans, subjects completed a questionnaire assessing comprehension of the story. The questionnaire comprised 27 context-independent and 12 context-dependent questions (39 questions in total). The resulting comprehension scores are reported as the proportion of correct answers (ranging 0–1) in the `participants.tsv` file and `scans.tsv` file for each subject. We recommend excluding the “Pretty Mouth and Green My Eyes” scans for subjects `sub-038` and `sub-105` (as specified in the `scan_exclude.json` file in the `code/` directory). The “Pretty Mouth and Green My Eyes” data have been reported in existing publications^108,203^. In the filename convention, “Pretty Mouth and Green My Eyes” is labeled using the task alias `prettymouth`.

### “Milky Way”

The “Milky Way” dataset was collected between March, 2013, and April, 2017, and comprised 53 participants (ages 18–34 years, mean age 21.7 ± 4.1 years, 27 reported female). The “Milky Way” story stimuli were written by an affiliate of the experimenters and read by a member of the Princeton Neuroscience Institute not affiliated with the experimenters’ lab. There are three versions of the “Milky Way” story capturing three different experimental conditions: (*a*) the original story (labeled `original`), which describes a man who visits a hypnotist to overcome his obsession with an ex-girlfriend, but instead becomes fixated on Milky Way candy bars; (*b*) an alternative version (labeled `vodka`) where sparse word substitutions yield a narrative in which a woman obsessed with an American Idol judge visits a psychic and becomes fixated on vodka; (*c*) a control version (labeled `synonyms`) with a similar number of word substitutions to the `vodka` version, but instead using synonyms of words in the original version, yielding a very similar narrative to the original. Relative to the original version, both the `vodka` and `synonyms` versions substituted on average 2.5 ± 1.7 words per sentence (34 ± 21% of words per sentence). All three versions were read by the same actor and each sentence of the recording was temporally aligned across versions. All three versions of the “Milky Way” story were 438 seconds (7.3 min; 292 TRs) long and began with 18 seconds of neutral introductory music followed by a 3 seconds of silence (21 TRs total), such that the story started at 0:21 and ended at 7:05, for a duration 404 s, with 13 seconds of silence at the end of the scan. The functional runs comprised 297 volumes (444.5 seconds). The transcript for the `original` version of the stimulus contains 1,059 words; 7 words (0.7%) were not successfully localized by the forced-alignment algorithm and 16 words (1.5%) were not found in the vocabulary. The `vodka` version contains 1058 words; 2 words (0.2%) were not successfully localized and 21 words (2.0%) were not found in the vocabulary. The `synonyms` version contains 1,066 words; 10 words (0.9%) were not successfully localized and 13 words (1.2%) were not found in the vocabulary.

The three versions of the stimuli were assigned to subjects according to a between-subject design, such that there were 18 subjects in each of the three groups, and each subject received only one version of the story. The group assignments are indicated in the `participants.tsv` file, and the `scans.tsv` file for each subject. The stimulus filename includes the version (`milkywayoriginal`, `milkywayvodka`, `milkywaysynonyms`) and is specified in the `events.tsv` file accompanying each run. The data corresponding to the `original` and `vodka` versions were collected between March and October, 2013, while the `synonyms` data were collected in March and April, 2017. Subjects completed a 28-item questionnaire assessing story comprehension following the scanning session. The comprehension scores for the `original` and `vodka` groups are reported as the proportion of correct answers (ranging 0–1) in the `participants.tsv` file and `scans.tsv` file for each subject. We recommend excluding “Milky Way” scans for subjects `sub-038`, `sub-105`, and `sub-123` (as specified in the `scan_exclude.json` file in the `code/` directory). The “Milky Way” data have been previously reported^196^. In the filename convention, “Milky Way” is labeled using the task alias `milkyway`.

### “Slumlord” and “Reach for the Stars One Small Step at a Time”

The “Slumlord” and “Reach for the Stars One Small Step at a Time” dataset was collected between May, 2013, and October, 2013, and comprised 18 participants (ages 18–27 years, mean age 21.0 ± 2.3 years, 8 reported female). The “Slumlord” story was told by Jack Hitt and recorded live at The Moth, a non-profit storytelling event, in New York City in 2006 (freely available at https://themoth.org/stories/slumlord). The “Reach for the Stars One Small Step at a Time” story was told by Richard Garriott and also recorded live at The Moth in New York City in 2010 (freely available at https://themoth.org/stories/reach-for-the-stars). The combined audio stimulus is 1,802 seconds (~30 minutes) long in total and begins with 22.5 seconds of music followed by 3 seconds of silence. The “Slumlord” story begins at approximately 0:25 relative to the beginning of the stimulus file, ends at 15:28, for a duration of 903 seconds (602 TRs), and is followed by 12 seconds of silence. After another 22 seconds of music starting at 15:40, the “Reach for the Stars One Small Step at a Time” story starts at 16:06 (965 seconds; relative to the beginning of the stimulus file), ends at 29:50, for a duration of 825 seconds (~550 TRs), and is followed by 12 seconds of silence. The stimulus file was started after 3 TRs (4.5 seconds) as indicated in the `events.tsv` files accompanying each scan. The scans were acquired with a variable number of trailing volumes following the stimulus across subjects, but can be truncated as researchers see fit (e.g. to 1205 TRs). The transcript for the combined “Slumlord” and “Reach for the Stars One Small Step at a Time” stimulus contains 5,344 words; 116 words (2.2%) were not successfully localized by the forced-alignment algorithm, and 57 words (1.1%) were not found in the vocabulary. The transcript for “Slumlord” contains 2,715 words; 65 words (2.4%) were not successfully localized and 25 words (0.9%) were not found in the vocabulary. The transcript for “Reach for the Stars One Small Step at a Time” contains 2,629 words; 51 words (1.9%) were not successfully localized and 32 words (1.2%) were not found in the vocabulary. We recommend excluding `sub-139` due to a truncated acquisition time (as specified in the `scan_exclude.json` file in the `code/` directory).

After the scans, each subject completed an assessment of their comprehension of the “Slumlord” story. To evaluate comprehension, subjects were presented with a written transcript of the “Slumlord” story with 77 blank segments, and were asked to fill in the omitted word or phrase for each blank (free response) to the best of their ability. The free responses were evaluated using Amazon Mechanical Turk (MTurk) crowdsourcing platform. MTurk workers rated the response to each omitted segment on a scale from 0–4 of relatedness to the correct response, where 0 indicates no response provided, 1 indicates an incorrect response unrelated to the correct response, and 4 indicates an exact match to the correct response. The resulting comprehension scores are reported as the proportion out of a perfect score of 4 (ranging 0–1) in the `participants.tsv` file and `scans.tsv` file for each subject. Comprehension scores for “Reach for the Stars One Small Step at a Time” are not provided. The “Slumlord” and “Reach for the Stars One Small Step at a Time” data have not been previously reported; however, a separate dataset not included in this release has been collected using the “Slumlord” story stimulus^204^. In the filename convention, the combined “Slumlord” and “Reach for the Stars One Small Step at a Time” data are labeled `slumlordreach`, and labeled `slumlord` and `reach` when analyzed separately.

### “It’s Not the Fall that Gets You”

The “It’s Not the Fall that Gets You” dataset was collected between May, 2013, and October, 2013, and comprised 56 participants (ages 18–29 years, mean age 21.0 ± 2.4 years, 31 reported female). The “It’s Not the Fall that Gets You” story was told by Andy Christie and recorded live at The Moth, a non-profit storyteller event, in New York City in 2009 (freely available at https://themoth.org/stories/its-not-the-fall-that-gets-you). In addition to the original story (labeled `intact`), two experimental manipulations of the story stimulus were created: (*a*) in one version, the story stimulus was temporally scrambled at a coarse level (labeled `longscram`); (*b*) in the other version, the story stimulus was temporally scrambled at a finer level (labeled `shortscram`). In both cases, the story was spliced into segments at the level of sentences prior to scrambling, and segment boundaries used for the `longscram` condition are a subset of the boundaries used for the `shortscram` condition. The boundaries used for scrambling occurred on multiples of 1.5 seconds to align with the TR during scanning acquisition. We recommend using only the intact version for studying narrative processing because the scrambled versions do not have a coherent narrative structure (by design). All three versions of the “It’s Not the Fall that Gets You” stimulus were 582 seconds (9.7 minutes) long and began with 22 seconds of neutral introductory music followed by 3 seconds of silence such that the story started at 0:25 (relative to the beginning of the stimulus file) and ended at 9:32 for a duration of 547 seconds (365 TRs), with 10 seconds of silence at the end of the stimulus. The stimulus file was started after 3 TRs (4.5 seconds) as indicated in the `events.tsv` files accompanying each scan. The functional scans comprised 400 TRs (600 seconds). The transcript for the `intact` stimulus contains 1,601 words total; 40 words (2.5%) were not successfully localized by the forced-alignment algorithm and 20 words (1.2%) were not found in the vocabulary.

The scrambled stimuli were assigned to subjects according to a mixed within- and between-subject design, such that subjects received the intact stimulus and either the longscram stimulus (23 participants) or the shortscram stimulus (24 participants). The group assignments are indicated in the `participants.tsv` file, and the `scans.tsv` file for each subject. Due to the mixed within- and between-subject design, the files are named with the full `notthefallintact`, `notthefalllongscram`, `notthefallshortscram` task labels. We recommend excluding the `intact` scans for `sub-317` and `sub-335`, the `longscram` scans for `sub-066` and `sub-335`, and the shortscram scan for `sub-333` (as specified in the `scan_exclude.json` file in the `code/` directory). The “It’s Not the Fall that Gets You” data have been recently reported^205^.

### “Merlin” and “Sherlock”

The “Merlin” and “Sherlock” datasets were collected between May, 2014, and March, 2015, and comprised 36 participants (ages 18–47 years, mean age 21.7 ± 4.7 years, 22 reported female). The “Merlin” and “Sherlock” stimuli were recorded while an experimental participant recalled events from previously viewed television episodes during fMRI scanning. Note that this spontaneous recollection task (during fMRI scanning) is cognitively distinct from reciting a well-rehearsed story to an audience, making this dataset different from others in this release. This release contains only the data for subjects listening to the auditory verbal recall; not data for the audiovisual stimuli or the speaker’s fMRI data^206^. The “Merlin” stimulus file is 915 seconds (9.25 minutes) long and began with 25 seconds of music followed by 4 seconds of silence such that the story started at 0:29 and ended at 15:15 for a duration of 886 seconds (591 TRs). The transcript for the “Merlin” stimulus contains 2,245 words; 111 words (4.9%) were not successfully localized by the forced-alignment algorithm and 13 words (0.6%) were not found in the vocabulary. The “Sherlock” stimulus file is 1,081 seconds (~18 minutes) long and began with 25 seconds of music followed by 4 seconds of silence such that the story started at 0:29 and ended at 18:01 for a duration of 1,052 seconds (702 TRs). The stimulus files for both stories were started after 3 TRs (4.5 seconds) as indicated in the `events.tsv` files accompanying each scan. The “Merlin” and “Sherlock” scans were acquired with a variable number of trailing volumes following the stimulus across subjects, but can be truncated as researchers see fit (e.g. to 614 and 724 TRs, respectively). The transcript for “Sherlock” contains 2681 words; 171 words (6.4%) were not successfully localized and 17 words (0.6%) were not found in the vocabulary. The word counts for “Merlin” and “Sherlock” may be slightly inflated due to the inclusion of disfluencies (e.g. “uh”) in transcribing the spontaneous verbal recall.

All 36 subjects listened to both the “Merlin” and “Sherlock” verbal recall auditory stimuli. However, 18 subjects viewed the “Merlin” audiovisual clip prior to listening to both verbal recall stimuli, while the other 18 subjects viewed the “Sherlock” audiovisual clip prior to listening to both verbal recall stimuli. In the “Merlin” dataset, the 18 subjects that viewed the “Sherlock” clip (and not the “Merlin” audiovisual clip) are labeled with the `naive` condition as they heard the “Merlin” verbal recall auditory-only stimulus without previously having seen the “Merlin” audiovisual clip. The other 18 subjects in the “Merlin” dataset viewed the “Merlin” audiovisual clip (and not the “Sherlock” clip) prior to listening to the “Merlin” verbal recall and are labeled with the `movie` condition. Similarly, in the “Sherlock” dataset, the 18 subjects that viewed the “Merlin” audiovisual clip (and not the “Sherlock” audiovisual clip) are labeled with the `naive` condition, while the 18 subjects that viewed the “Sherlock” audiovisual clip (and not the “Merlin” audiovisual clip) are labeled with the `movie` condition. These condition labels are indicated in both the `participants.tsv` file and the `scans.tsv` files for each subject. Following the scans, each subject performed a free recall task to assess memory of the story. Subjects were asked to write down events from the story in as much detail as possible with no time limit. The quality of comprehension for the free recall text was evaluated by three independent raters on a scale of 1–10. The resulting comprehension scores are reported as the sum across raters normalized by the perfect score (range 0–1) in the `participants.tsv` file and `scans.tsv` file for each subject. Comprehension scores are only provided for the `naive` condition. We recommend excluding the “Merlin” scan for subject `sub-158` and the “Sherlock” scan for `sub-139` (as specified in the `scan_exclude.json` file in the `code/` directory. The “Merlin” and “Sherlock” data have been previously reported^206^. In the filename convention, “Merlin” is labeled `merlin` and “Sherlock” is labeled `sherlock`.

### “Schema”

The “Schema” dataset was collected between August, 2015, and March, 2016, and comprised 31 participants (ages 18–35 years, mean age 23.7 ± 4.8 years, 13 reported female). The “Schema” dataset comprised eight brief (~3-minute) auditory narratives based on popular television series and movies featuring restaurants and airport scenes: *The Big Bang Theory, Friends, How I Met Your Mother, My Cousin Vinnny, The Santa Clause, Shame, Seinfeld, Up in the Air*; labeled `bigbang`, `friends`, `himym`, `vinny`, `santa`, `shame`, `seinfeld`, `upintheair`, respectively. These labels are indicated in the `events.tsv` accompanying each scanner run, and the corresponding stimuli (e.g. `bigbang_audio.wav`). Two of the eight stories were presented in each of four runs. Each run had a fixed presentation order of a fixed set of stories, but the presentation order of the four runs was counterbalanced across subjects. The functional scans were acquired with a variable number of trailing volumes across subjects, and in each run two audiovisual film clips (not described in this data release) were also presented; the auditory stories must be spliced from the full runs according to the `events.tsv` file. Overall, the transcripts for the auditory story stimuli in the “Schema” dataset contain 3,788 words; 20 words (0.5%) were not successfully localized by the forced-alignment algorithm and 69 words (1.8%) were not found in the vocabulary. The “Schema” data were previously reported^115^. In the filename convention, the “Schema” data are labeled using the task alias schema.

### “Shapes”

The “Shapes” dataset was collected between May, 2015, and July, 2016, and comprised 59 participants (ages 18–35 years, mean age 23.0 ± 4.5 years, 42 reported female). The “Shapes” dataset includes two related auditory story stimuli describing a 7-minute original animation called “When Heider Met Simmel” (copyright Yale University). The movie uses two-dimensional geometric shapes to tell the story of a boy who dreams about a monster, and was inspired by Heider and Simmel^207^. The movie itself is available for download at https://www.headspacestudios.org. The two verbal descriptions based on the movie used as auditory stimuli in the “Shapes” dataset were: (*a*) a purely physical description of the animation (labeled `shapesphysical`), and (*b*) a social description of the animation intended to convey intentionality in the animated shapes (labeled `shapessocial`). Note that the physical description story (`shapesphysical`) is different from other stimuli in this release in that it describes the movements of geometrical shapes without any reference to the narrative embedded in the animation. Both stimulus files are 458 seconds (7.6 minutes) long and began with a 37-second introductory movie (only the audio channel is provided in the stimulus files) followed by 7 seconds of silence such that the story started at 0:45 and ended at roughly 7:32 for a duration of ~408 seconds (~272 TRs) and ended with ~5 seconds of silence. The stimulus files for both stories were started after 3 TRs (4.5 seconds) as indicated in the `events.tsv` files accompanying each scan. The functional scans were acquired with a variable number of trailing volumes following the stimulus across subjects, but can be truncated as researchers see fit (e.g. to 309 TRs). The transcript for the `shapesphysical` stimulus contains 951 words; 9 words (0.9%) were not successfully localized by the forced-alignment algorithm and 25 words (2.6%) were not found in the vocabulary. The transcript for the `shapessocial` stimulus contains 910 words; 6 words (0.7%) were not successfully localized and 14 words (1.5%) were not found in the vocabulary.

Each subject received both the auditory physical description of the animation (`shapesphysical`), the auditory social description of the animation (`shapessocial`), and the audiovisual animation itself (not included in the current data release) in three separate runs. Run order was counterbalanced across participants, meaning that some participants viewed the animation before hearing the auditory description. The run order for each subject (e.g. `physical-social-movie`) is specified in the `participants.tsv` file and the subject-specific `scans.tsv` files. Immediately following the story scan, subjects performed a free recall task in the scanner (the fMRI data during recall are not provided in this release). They were asked to describe the story in as much detail as possible using their own words. The quality of the recall was evaluated by an independent rater naive to the purpose of the experiment on a scale of 1–10. The resulting comprehension scores are reported normalized by 10 (range 0–1) and reported in the `participants.tsv` file and `scans.tsv` file for each subject. Comprehension scores are only provided for scans where the subject first listened to the story stimulus, prior to hearing the other version of the story or viewing the audiovisual stimulus. We recommend excluding the `shapessocial` scan for subject `sub-238` (as specified in the `scan_exclude.json` file in the `code/` directory. The “Shapes” data were previously reported^208^.

### “The 21st Year”

The “The 21st Year” dataset was collected between February, 2016, and January, 2018, and comprised 25 participants (ages 18–41 years, mean age 22.6 ± 4.7 years, 13 reported female). The story stimulus was written and read aloud by author Christina Lazaridi, and includes two seemingly unrelated storylines relayed in blocks of prose that eventually fuse into a single storyline. The stimulus file is 3,374 seconds (56.2 minutes) long and began with 18 seconds of introductory music followed by 3 seconds such that the story started at 0:21 and ended at 55:59 for a duration of 3,338 seconds (2,226 TRs) followed by 15 seconds of silence. The functional scans comprised 2,249 volumes (3,373.5 seconds). The story stimulus contains 8,267 words; 88 words (1.1%) were not successfully localized by the forced-alignment algorithm and 143 words (1.7%) were not found in the vocabulary. Data associated with “The 21st Year” have been previously reported^209^. In the filename convention, data for “The 21st Year” are labeled using the task alias `21styear`.

### “Pie Man (PNI)” and “Running from the Bronx”

The “Pie Man (PNI)” and “Running from the Bronx” data were collected between May and September, 2018, and comprised 47 participants (ages 18–53 years, mean age 23.3 ± 7.4 years, 33 reported female). The “Pie Man (PNI)” and “Running from the Bronx” story stimuli were told by Jim O’Grady while undergoing fMRI scans at the Princeton Neuroscience Institute (PNI). The spoken story was recorded using a noise cancelling microphone mounted on the head coil and scanner noise was minimized using Audacity. The “Pie Man (PNI)” stimulus file is 415 seconds (~7 minutes) long and begins with a period of silence such that the story starts at :09 and ends at 6:49 for a duration of 400 seconds (267 TRs) followed by 7 seconds of silence. The “Pie Man (PNI)” stimulus conveys the same general story as the original “Pie Man” stimulus but with differences in timing and delivery (as well as poorer audio quality due to being recorded during scanning). The “Running from the Bronx” stimulus file is 561 seconds (9.4 minutes) long and begins with a period of silence such that the story starts at 0:15 and ends at 9:11 for a duration of 536 seconds (358 TRs) followed by 10 seconds of silence. The “Pie Man (PNI)” and “Running from the Bronx” functional scans comprised 294 TRs (441 seconds) and 390 TRs (585 seconds), respectively. Both stimulus files were started after 8 TRs (12 seconds) as indicated in the accompanying `events.tsv` files. The program card on the scanner computer was organized according to the ReproIn convention^210^ to facilitate automated conversion to BIDS format using HeuDiConv 0.5.dev1^211^. The transcript for the “Pie Man (PNI)” stimulus contains 992 words; 7 words (0.7%) were not successfully localized by the forced-alignment algorithm and 19 words (1.9%) were not found in the vocabulary. The transcript for “Running from the Bronx” contains 1,379 words; 23 words (1.7%) were not successfully localized and 19 words (1.4%) were not found in the vocabulary.

At the end of the scanning session, the subjects completed separate questionnaires for each story to evaluate comprehension. Each questionnaire comprised 30 multiple choice and fill-in-the-blank questions testing memory and understanding of the story. The resulting comprehension scores are reported as the proportion of correct answers (ranging 0–1) in the `participants.tsv` file and the `scans.tsv` file for each subject. Data for both stories have been previously reported^165,212^. The “Pie Man (PNI)” and “Running from the Bronx” data are labeled with the piemanpni and bronx task aliases in the filename convention. The “Pie Man (PNI)” and “Running from the Bronx” data were collected in conjunction with the “I Knew You Were Black” and “The Man Who Forgot Ray Bradbury” data and share the same samples of subjects.

### “I Knew You Were Black” and “The Man Who Forgot Ray Bradbury”

The “I Knew You Were Black” and “The Man Who Forgot Ray Bradbury” data were collected by between May and September, 2018, and comprised the same sample of subjects from “Pie Man (PNI)” and “Running from the Bronx” data (47 participants, ages 18–53 years, mean age 23.3 ± 7.4 years, 33 reported female). Unlike the “Pie Man (PNI)” and “Running from the Bronx” story stimuli, the “I Knew You Were Black” and “The Man Who Forgot Ray Bradbury” stimuli are professionally recorded with high audio quality. The “I Knew You Were Black” story was told by Carol Daniel and recorded live at The Moth, a non-profit storytelling event, in New York City in 2018 (freely available at https://themoth.org/stories/i-knew-you-were-black). The “I Knew You Were Black” story is 800 seconds (13.3 minutes, 534 TRs) long and occupies the entire stimulus file. The “I Knew You Were Black” functional scans comprised 550 TRs (825 seconds). The transcript for the “I Knew You Were Black” stimulus contains 1,544 words; 5 words (0.3%) were not successfully localized by the forced-alignment algorithm and 4 words (0.3%) were not found in the vocabulary. “The Man Who Forgot Ray Bradbury” was written and read aloud by author Neil Gaiman at the Aladdin Theater in Portland, OR, in 2011 (freely available at https://soundcloud.com/neilgaiman/the-man-who-forgot-ray-bradbury). The “The Man Who Forgot Ray Bradbury” audio stimulus file is 837 seconds (~14 minutes, 558 TRs) long and occupies the entire stimulus file. The “The Man Who Forgot Ray Bradbury” functional scans comprised 574 TRs (861 seconds). The transcript for “The Man Who Forgot Ray Bradbury” contains 2,135 words; 16 words (0.7%) were not successfully localized and 29 words (1.4%) were not found in the vocabulary. For both datasets, the audio stimulus was prepended by 8 TRs (12 seconds) and followed by 8 TRs (12 seconds) of silence. The program card on the scanner computer was organized according to the ReproIn convention^210^ to facilitate automated conversion to BIDS format using HeuDiConv 0.5.dev1^211^.

Similarly to the “Pie Man (PNI)” and “Running from the Bronx” stories, the subjects completed separate questionnaires for each story to evaluate comprehension after scanning. Each questionnaire comprised 25 multiple choice and fill-in-the-blank questions testing memory and understanding of the story. The resulting comprehension scores are reported as the proportion of correct answers (ranging 0–1) in the `participants.tsv` file and the `scans.tsv` file for each subject. Data for both stories have been previously reported^165,212^. The “I Knew You Were Black” and “The Man Who Forgot Ray Bradbury” data are labeled with the black and forgot task aliases in the filename convention.

## Data records

The core, unprocessed NIfTI-formatted MRI data with accompanying metadata and stimuli are publicly available on OpenNeuro: https://openneuro.org/datasets/ds002345 (https://doi.org/10.18112/openneuro.ds002345.v1.1.4). All data and derivatives are hosted at the International Neuroimaging Data-sharing Initiative (INDI)^213^ (RRID:SCR_00536) via the Neuroimaging Informatics Tools and Resources Clearinghouse (NITRC)^214^ (https://www.nitrc.org/; RRID:SCR_003430): http://fcon_1000.projects.nitrc.org/indi/narratives/ (https://doi.org/10.15387/fcp_indi.retro.Narratives). The full data collection is available via the DataLad data distribution (RRID:SCR_019089): http://datasets.datalad.org/?dir=/labs/hasson/narratives.

Data and derivatives have been organized according to the machine-readable Brain Imaging Data Structure (BIDS) 1.2.1^85^, which follows the FAIR principles for making data findable, accessible, interoperable, and reusable^215^. A detailed description of the BIDS specification can be found at http://bids.neuroimaging.io/. Organizing the data according to the BIDS convention facilitates future research by enabling the use of automated BIDS-compliant software (BIDS Apps)^216^. Briefly, files and metadata are labeled using key-value pairs where key and value are separated by a hyphen, while key-value pairs are separated by underscores. Each subject is identified by an anonymized numerical identifier (e.g. `sub-001`). Each subject participated in one or more story-listening scans indicated by the task alias for the story (e.g. `task-pieman`). Subject identifiers are conserved across datasets; subjects contributing to multiple datasets are indexed by the same identifier (e.g. `sub-001` contributes to both the “Pie Man” and “Tunnel Under the World” datasets).

The top-level BIDS-formatted `narratives/` directory contains a tabular `participants.tsv` file where each row corresponds to a subject and includes demographic variables (`age`, `sex`), the stories that subject received (`task`), as well as experimental manipulations (`condition`) and comprehension scores where available (`comprehension`). Cases of omitted, missing, or inapplicable data are indicated by “n/a” according to the BIDS convention. The top-level BIDS directory also contains the following: (*a*) a `dataset_description.json` containing top-level metadata, including licensing, authors, acknowledgments, and funding sources; (*b*) a `code/` directory containing scripts used for analyzing the data; (*c*) a `stimuli/` directory containing the audio stimulus files (e.g. `pieman_audio.wav`), transcripts (e.g. `pieman_transcript.txt`), and a `gentle/` directory containing time-stamped transcripts created using Gentle (i.e.; described in “Stimuli” in the “Methods” section); (*d*) a `derivatives/` directory containing all MRI derivatives (described in more detail in the “MRI preprocessing” and “Technical validation” sections); and (*e*) descriptive `README` and `CHANGES` files.

BIDS-formatted MRI data for each subject are contained in separate directories named according to the subject identifiers. Within each subject’s directory, data are segregated into anatomical data (the `anat/` directory) and functional data (the `func/` directory). All MRI data are stored as gzipped (compressed) NIfTI-1 images^217^. NIfTI images and the relevant metadata were reconstructed from the original Digital Imaging and Communications in Medicine (DICOM) images using dcm2niix 1.0.20180518^218^. In instances where multiple scanner runs were collected for the same imaging modality or story in the same subject, the files are differentiated using a run label (e.g. `run-1`, `run-2`). All anatomical and functional MRI data files are accompanied by sidecar JSON files containing MRI acquisition parameters in compliance with the COBIDAS report^132^. Identifying metadata (e.g. name, birth date, acquisition date and time) have been omitted, and facial features have been removed from anatomical images using the automated de-facing software pydeface 2.0.0^133^. All MRI data and metadata are released under the Creative Commons CC0 license (https://creativecommons.org/), which allows for free reuse without restriction.

All data and derivatives are version-controlled using DataLad^87,88^ (https://www.datalad.org/; RRID:SCR_003931). DataLad uses git (https://git-scm.com/; RRID:SCR_003932) and git-annex (https://git-annex.branchable.com/; RRID:SCR_019087) to track changes made to the data, providing a transparent history of the dataset. Each stage of the dataset’s evolution is indexed by a unique commit hash and can be interactively reinstantated. Major subdivisions of the data collection, such as the `code/`, `stimuli/`, and derivative directories, are encapsulated as “subdatasets” nested within the top-level BIDS dataset. These subdatasets can be independently manipulated and have their own standalone version history. For a primer on using DataLad, we recommend the DataLad Handbook^219^ (http://handbook.datalad.org/).

## Technical validation

### Image quality metrics

To provide an initial assessment of the data, we used MRIQC 0.15.1^220,221^ to derive a variety of image quality metrics (IQMs; via the `run_mriqc.sh` and `run_mriqc_group.sh` scripts in the `code/` directory). MRIQC was deployed using Singularity^139^. The MRIQC Singularity image can be built from Docker Hub (https://hub.docker.com/r/poldracklab/mriqc/; e.g. `singularity build mriqc-0.15.1.simg docker://poldracklab/mriqc:0.15.1`). The MRIQC outputs can be found in the `mriqc/` directory in `derivatives/` available via the DataLad release, and includes dozens of IQMs per scan as well as a summary across the entire dataset. These IQMs can be visualized in a browser (e.g. `group_bold.html`) and are aggregated in tabular format across the entire data collection (e.g. `group_bold.tsv`). Here we do not exclude any subjects based on IQMs, but subsequent researchers can use the available IQMs to exclude scans as they see fit.

We briefly report three common metrics for functional images summarizing head motion, intrinsic spatial smoothness, and temporal signal quality (Fig. 2). To summarize head motion, we computed the mean framewise displacement (FD)^161,222^ for each functional scan across all subjects and story stimuli (Fig. 2a). The median framewise displacement across scans for all subjects and story stimuli was 0.132 mm (*SD* = 0.058 mm). Although this indicates relatively low head motion, some subjects have notably high motion; researchers may opt to censor time points with high FD (e.g. greater 0.5 mm) or omit entire scans with high overall FD as they see fit. To quantify the intrinsic spatial smoothness of the raw functional data, we used `3dFWHMx` in AFNI 19.3.0, which computes the ratio of variance of first differences across voxels to global variance across the image and expresses smoothness as the full width at half-maximum (FWHM) of a 2D Gaussian filter^223^. Smoothness was computed in each subject’s native brain space within an anatomically-defined brain mask, and the functional time series were temporally detrended prior to smoothness estimation (using the `get_fwhm.py` script in the `code/` directory). Spatial smoothness varied considerably across datasets due to different acquisition parameters used with different scanner models (Fig. 2b). For example, scans acquired using older pulse sequences without multiband acceleration are smoothest in the slice acquisition (*z*-) axis, whereas newer scans acquired using multiband acceleration have higher overall spatial resolution (resulting in lower smoothness), and are smoothest in anterior–posterior phase-encoding (*y*-) axis. Finally, we computed the temporal signal-to-noise ratio (tSNR) as a measure of functional signal quality using MRIQC. A tSNR map was constructed for each scan by computing the voxelwise average of the signal magnitude over time divided by the standard deviation of the signal over time in each subject’s native space following realignment using `3dvolreg` and brain extraction using `3dSkullStrip` in AFNI^224^; the median of this tSNR map was used to summarize tSNR across voxels for a given scan (Fig. 2c). The median tSNR across all subjects and story stimuli was 50.772 (*SD* = 13.854). We also computed vertex-wise tSNR in fsaverage6 space following preprocessing with fMRIPrep to visualize the spatial distribution of tSNR across cortex (Fig. 2d; using the `get_tsnr.py` script in the `code/` directory). Low tSNR in orbitofrontal and anterior medio-temporal cortex reflects signal dropout due to the proximity of sinuses and ear canals^225^.

**Fig. 2.**
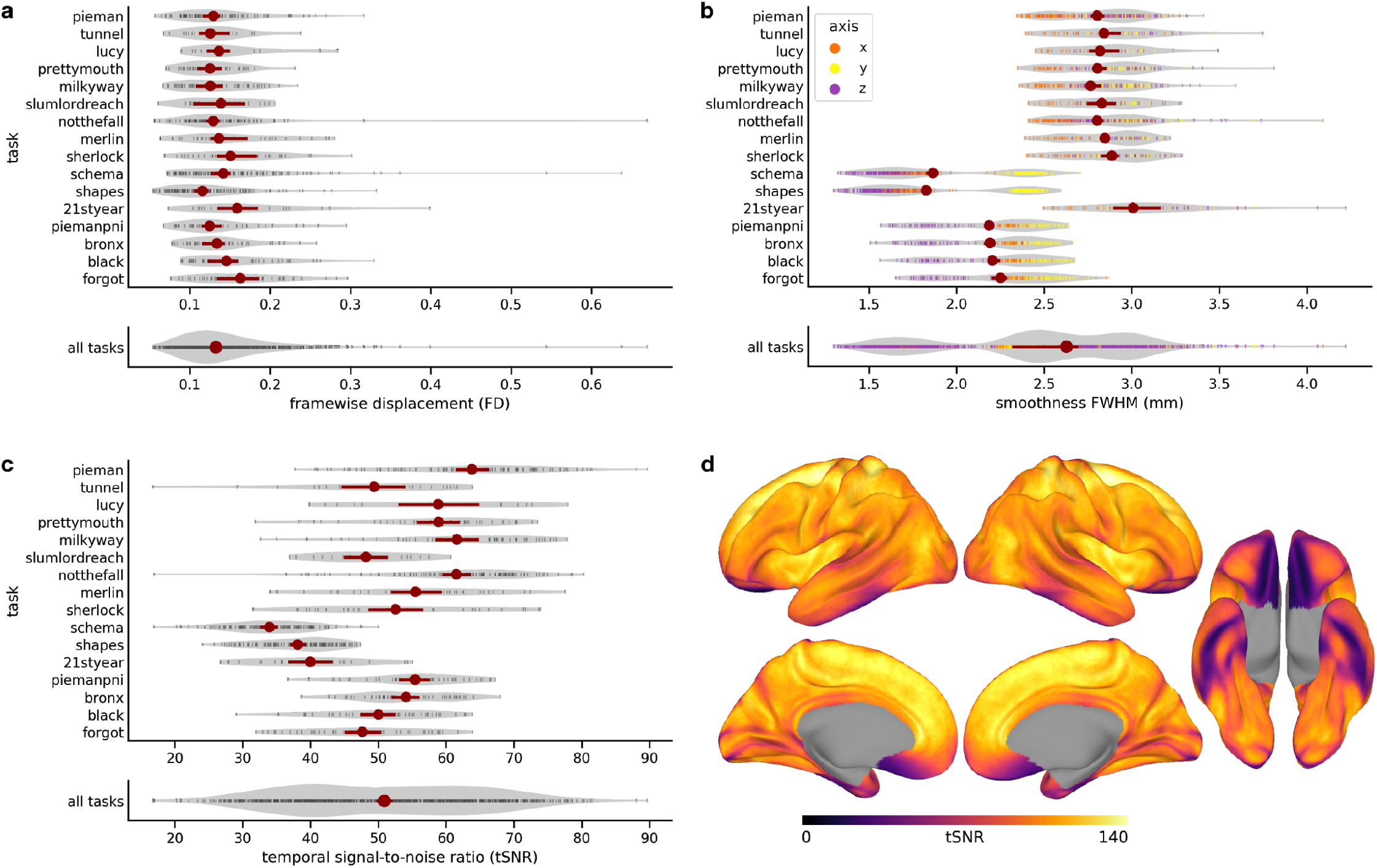
Data quality summarized in terms of head motion, spatial smoothness, and temporal signal-to-noise ratio (tSNR). (**a**) Median framewise displacement (FD) summarizes head motion. Each black tick mark represents the median FD (*x*-axis) for a given scanning run and subject for each task (i.e. story; *y*-axis). Violin plots illustrate the distribution of median FDs across all runs and subjects for a given task. The red markers indicate the median FD across all runs for a given task, and the red error bars indicate the 95% bootstrap confidence interval for the median based on randomly sampling runs with replacement. At bottom, FD is summarized across all tasks. (**b**) Spatial smoothness is summarized using AFNI’s FWHM smoothness for each scanning run across each task. Spatial smoothness was computed in volumetric subject-specific functional (EPI) space using subject-specific whole-brain masks with minimal preprocessing (realignment and susceptibility distortion in fMRIPrep, detrending in AFNI). Each tick mark represents the spatial smoothness for a given run in a given acquisition axis (orange: *x*-axis, i.e. left–right; yellow: *y*-axis, i.e. anterior–posterior; red: *z*-axis, i.e. inferior–superior). Violin plots capture the distribution of smoothness estimates for *x*-, *y*-, and *z*-axes across all runs and subjects in a given task. The red markers indicate the median combined (i.e. geometric mean across *x*-, *y*-, and *z*-axes) smoothness across all runs and subjects for a given task (red error bars indicate 95% bootstrap confidence interval of median). At bottom, smoothness is summarized across all tasks. The multiple lobes of the distribution reflect differing acquisition parameters used with the Skyra and Prisma scanner models. (**c**) tSNR maps were constructed by computing the voxelwise mean signal over time divided by the standard deviation over time. Each black tick mark represents the median of the tSNR map (*x*-axis) for a given scanner run and subject for each task (*y*-axis). Violin plots reflect the distribution of tSNR values across all runs and subjects for a given task. The red markers indicate the mean tSNR across all runs for a given task (red error bars indicate 95% bootstrap confidence interval of mean). At bottom, the tSNR is summarized across all tasks. The two lobes of the distribution reflect older pulse sequences with larger voxels (and higher tSNR) and newer multiband pulse sequences with smaller voxels (and lower tSNR). See the `plot_qc.py` script in the `code/` directory for details. (**d**) Distribution of tSNR across cortex. The color bar reflects the median vertex-wise tSNR. Note that unlike panel **c**, here tSNR was computed in fsaverage6 space for vertex-wise summarization and visualization, which may inflate the tSNR values.

### Intersubject correlation

To measure the reliability of stimulus-evoked responses across subjects, we performed an intersubject correlation (ISC) analysis^103,104,226^. For each story stimulus, we computed ISC using the leave-one-out approach: for each subject, we averaged the response time series for the remaining subjects, then correlated this mean response time series for a given voxel or brain area with the response time series for the left-out subject in that voxel or brain area. Note that this approach yields higher correlation values than computing ISC between pairs of subjects. In large samples of subjects, leave-one-out ISC provides an estimate of the upper bound of variance that can be accounted for by between-subject models predicting voxel- or region-wise response time series (i.e. noise ceiling)^104,227^; that is, the mean data from other subjects serves as a surrogate for the optimal model that generalizes across subjects (within-subject models, however, may capture additional variance not shared across subjects). ISC was computed only on functional volumes corresponding to presentation of the story stimulus, and we trimmed the first 6 TRs (9 seconds) after story onset to reduce the impact of stimulus onset transients. All ISC values were Fisher z-transformed prior to averaging, and the mean was inverse Fisher *z*-transformed for visualization^228^. ISC analysis was implemented using the Brain Imaging Analysis Kit (BrainIAK)^229–231^ (RRID:SCR_014824).

ISCs were computed separately for different experimental groups or manipulations; for example, ISCs were computed separately for the “affair” and “paranoid” contextual manipulation conditions for “Pretty Mouth and Green My Eyes.” On the other hand, ISCs were computed as a single group for datasets where the only experimental manipulation was naively listening to the story stimulus versus listening to the story stimulus after viewing the audiovisual movie (“Merlin,” “Sherlock,” and “Shapes”). ISCs were computed separately for the “Slumlord” and “Reach for the Stars One Small Step at a Time” stories despite being presented in the same scanning run. The brief “Schema” stories and the scrambled versions of “It’s Not the Fall that Gets You” were excluded from ISC analysis for the sake of simplicity.

To assess the temporal synchronization of responses to the story stimuli, we computed ISC in an early auditory cortex (EAC) region of interest (ROI). The left- and right-hemisphere EAC ROIs were defined on the cortical surface in *fsaverage6* space as combination of five areas (A1, MBelt, LBelt, PBelt, RI) from a multimodal cortical parcellation^232,233^ (see the `roi_masks.py` script in the `code/` directory). The non-smoothed response time series following preprocessing with fMRIPrep and confound regression with AFNI were averaged across vertices in the left and right EAC ROIs (using the `roi_average.py` and `roi_regression.py` scripts in the `code/` directory). We used two measures of temporal alignment in left EAC as quality-control criteria for excluding scans. First, we computed leave-one-out ISCs and set an exclusion cut-off at ISC < .10 in left EAC (Fig. 3a; using the `roi_isc.py` script in the `code/` directory). Second, we computed leave-one-out ISC at temporal lags ranging from −45 to 45 seconds; for each left-out subject, we computed the correlation between the mean response time series for the remaining subjects and the left-out subject’s response time series at 61 lags ranging from −30 to 30 TRs (Fig 3b; using the `roi_lags.py` script in the `code/` directory). We then set an exclusion cut-off at a peak ISC at a lag > ±3 TRs (4.5 seconds) to accommodate some inter-individual variability in the hemodynamic response^234,235^. Both of these criteria are intended to exclude scans with poor entrainment to the stimulus (e.g. due to an error in acquisition), and yield a partly overlapping group of scans for exclusion. Overall, the combined criteria resulted in excluding 27 scans from 19 subjects, or 3.0% of the 891 total scans. We do not exclude these subjects from the dataset entirely because some analyses (e.g. comparing methods for mitigating artefacts) may in fact benefit from the inclusion of lower-quality scans. Rather, we provide a `scan_exclude.json` file listing scans to exclude (and a convenience function `exclude_scans.py`) in the `code/` directory so that researchers can easily exclude scans flagged as poor quality according to the criteria above. Note that some studies^205^ temporally shift response time series to ensure that peak lags match across subjects; here, we do not shift or edit the scans, but provide the lagged ISCs for all scans (`group_roi-EAC_lags.json` in `derivatives/afni-nosmooth/`) so that researchers can apply temporal shifting as they see fit. Ultimately, mean ISC in left and right EAC across all scans was .549 (*SD* = .173) and .493 (*SD* = .179), respectively. This analysis also revealed qualitatively higher ISC in left EAC than right EAC across several stories (Fig. 3a). This may reflect a left-lateralized preference for speech sounds observed in previous studies^10,236,237^.

**Fig. 3.**
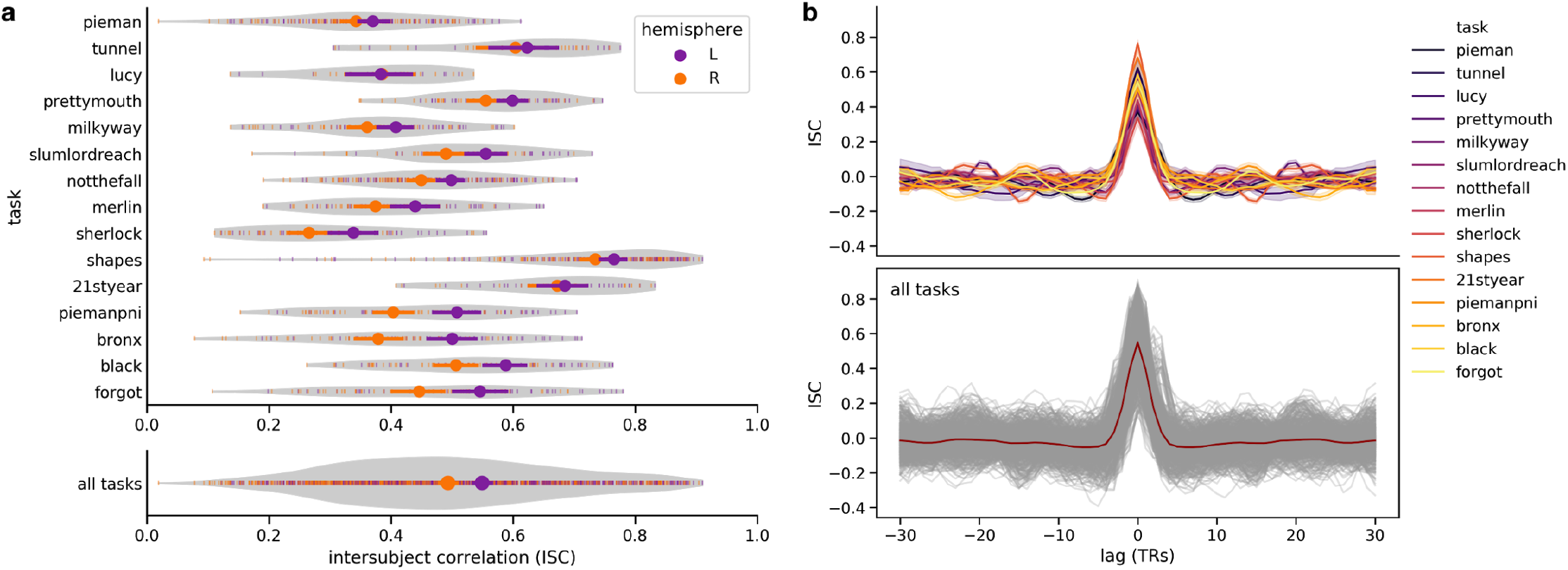
Intersubject correlation (ISC) in early auditory cortex (EAC). (**a**) ISC in left and right EAC across all subjects and tasks (i.e. stories). Each tick mark represents the leave-one-out ISC (*x*-axis) for a left-out subject in either left- (purple) or right- (orange) hemisphere EAC across tasks (*y*-axis). Violin plots illustrate the distribution of leave-one-out ISC across all left-out subjects for each task. The circular markers indicate the mean ISC across left-out subjects for each task and hemisphere (error bars indicate the 95% bootstrap confidence interval of the mean). At bottom, ISC for left and right EAC is summarized across all tasks. (**b**) Lagged ISC captures temporal synchronization in left EAC. Leave-one-out ISC (*y*-axis) was computed for left EAC at 61 lags ranging from −30 to +30 TRs (−45 to 45 seconds; *x*-axis). In the upper plot, each line represents the mean ISC across lags for each task. At bottom, all lagged ISCs are visualized in a single plot where each left-out subject corresponds to a semi-transparent gray line; the red line reflects the average lagged ISC across all subjects and tasks. See the `plot_isc.py` script in the `code/` directory for details.

Finally, we visualized the cortical distribution of ISCs (Fig. 4). Leave-one-out ISCs were computed on vertex-wise response time series in *fsaverage6* surface space following preprocessing using fMRIPrep, spatial smoothing to 6 mm, and confound regression using AFNI (via the `run_isc.py` script in the `code/` directory). ISCs were computed separately for each story stimulus (and experimental group or condition, as described above), then averaged across all stories to construct a summary ISC map (Fig. 4a). Here we are not interested in statistical testing and therefore arbitrarily threshold these ISC maps at a mean correlation of .10 for visualization; we refer researchers to other work for a detailed statistical treatment^238–240^. Cortical areas with high ISC during story-listening were largely bilateral, spanning temporal and frontal areas reported in the fMRI literature on language, and extending into default-mode network (DMN) areas such as the angular gyrus and precuneus. Cortical ISC maps are strikingly similar across stories with considerable variability in sample size, duration, and content (Fig. 4b). We emphasize that a wide variety of stimulus content can drive cortical ISCs, and that decomposing the stimuli according to different models will ultimately provide a window onto the linguistic and narrative features encoded across cortex.

**Fig. 4.**
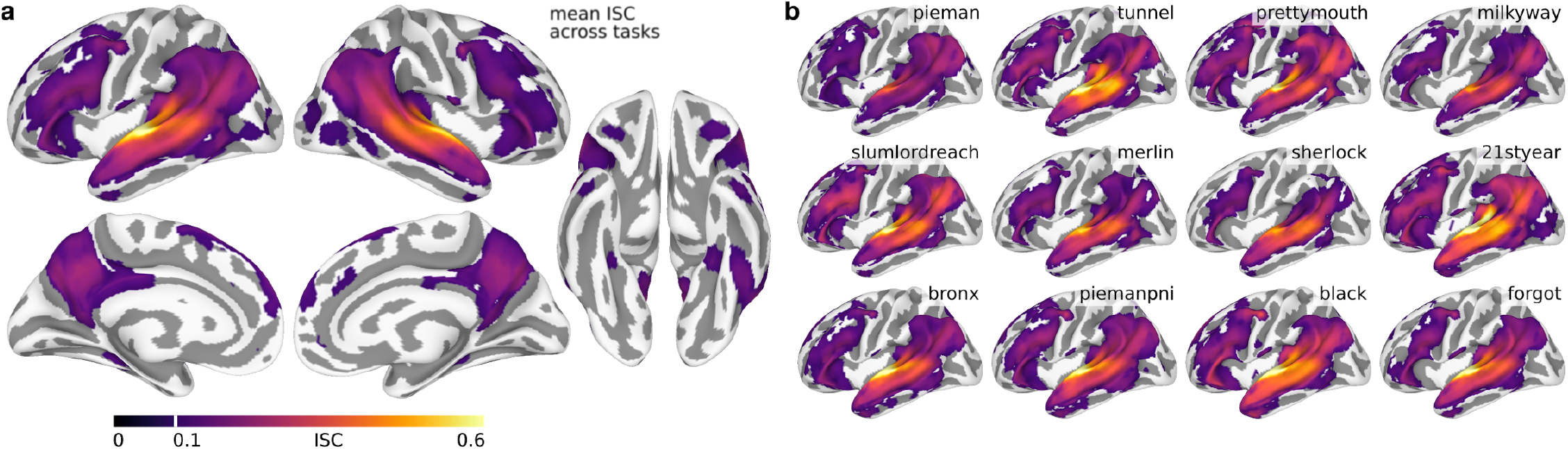
Intersubject correlations across cortex. (**a**) The mean vertex-wise leave-one-out ISC was computed across all subjects and stories. The color bar reflects the mean leave-one-out ISC and ranges from 0 to .60. ISC maps were arbitrarily thresholded at ISC = .10. (**b**) Mean ISC maps for 12 representative stories plotted according to the same color map and thresholded at ISC = .10. Where applicable (e.g. “Pretty Mouth and Green My Eyes”), ISCs were computed separately for different experimental groups or conditions, then averaged; ISCs were also computed separately for “Slumlord” and “Reach for the Stars One Small Step at a Time,” then averaged.

## Usage notes

The MRI data are organized according to the BIDS standard, and are therefore well-suited for automated processing using BIDS Apps (e.g. MRIQC, fMRIPrep, FitLins^241^; https://bids-apps.neuroimaging.io/), the Brainlife cloud platform (https://brainlife.io/), and Neuroscout^242^ (https://neuroscout.org/). The full data collection can be easily “installed” using DataLad (`datalad install -r ///labs/hasson/narratives`). This clones the file hierarchy without downloading any large content files, which can be installed selectively as needed (`datalad get`). Particular subsets of the full dataset can be retrieved using `datalad get` with shell wildcards. For example, particular stories can be retrieved using, e.g. `datalad get sub-*/func/*task-pieman*`; or derivatives in a particular output space can be retrieved using, e.g. `datalad get derivatives/afni-nosmooth/sub-*/func/*task-pieman*space-fsaverage6*`. For convenience, we also include the `task_meta.json` file in the code directory, which contains a dictionary mapping from each task (i.e. story) to subjects and filenames belonging to that task. We also recommend the Python-based PyBIDS tool for querying and manipulating BIDS data^243^.

## Code availability

All code used for aggregating and analyzing the data is version-controlled and publicly available via the associated GitHub repository (https://github.com/snastase/narratives) and the `code/` directory in the top-level BIDS directory distributed via DataLad (https://datasets.datalad.org/?dir=/labs/hasson/narratives). The GitHub repository contains both scripts used to prepare the data for sharing (`staging/`) and scripts used to analyze the BIDS-formatted data (`code/`). See Table S1 for a brief description of the scripts used to process the “Narratives” data.

## Acknowledgments

We thank Leigh Nystrom, Mark Pinsk, Garrett McGrath, and the administrative staff at the Scully Center for the Neuroscience of Mind and Behavior and the Princeton Neuroscience Institute, as well as Elizabeth McDevitt, Anne Mennen, and members of Pygers support group. We thank Franklin Feingold for assistance in data sharing, as well as Chris Gorgolewski, Tal Yarkoni, Satrajit S. Ghosh, Avital Hahamy, Mohamed Amer, Indranil Sur, Xiao Lin, and Ajay Divarakian for helpful feedback on the data and analysis. This work was supported by the National Institutes of Health (NIH) grants R01-MH094480 (U.H.), DP1-HD091948 (U.H.), R01-MH112566 (U.H.), R01-MH112357 (K.A.N., U.H), T32-MH065214 (K.A.N), by the Defense Advanced Research Projects Agency (DARPA) Brain-to-Brain Seedling contract number FA8750-18-C-0213 (U.H.), and by the Intel Corporation. The views, opinions, and/or conclusions contained in this paper are those of the authors and should not be interpreted as representing the official views or policies, either expressed or implied of the NIH, DARPA, or Intel.

## Author contributions

S.A.N. converted the data to BIDS format, analyzed the data, and wrote the paper. S.A.N., Y.F.L., L.H., H.H., A.Z., and E.M. curated the data. J.C., C.J.H., E. S., M.A.C., M.R. and U.H. designed and collected the “Pie Man” dataset. O.L. and U.H. designed and collected the “Tunnel Under the World” and “Lucy” datasets. Y.Y. and U.H. designed and collected the “Pretty Mouth and Green My Eyes” and “Milky Way”datasets. J.C., M.R., M.C. and U.H. designed and collected the “Slumlord”, “Reach for the Stars One Small Step at a Time”, and “It’s Not the Fall that Gets You” datasets. A.Z., J.C., Y.C.L., K.A.N., and U.H. designed and collected the “Merlin” and “Sherlock” datasets. C.B., U.H., and K.A.N. designed and collected the “Schema” dataset. M.N. and U.H. designed and collected the “Shapes” dataset. C.H.C.C., Y.Y., K.A.N., and U.H. designed and collected the “The 21st Year” dataset. S.N., P.P.B., and U.H. designed and collected the “Pie Man”, “Running from the Bronx”, “I Knew You Were Black”, and “The Man Who Forgot Ray Bradbury” datasets. G.C. and A.G. provided technical assistance in stimulus transcription. Y.O.H. provided technical assistance with DataLad. U.H., K.A.N., and Y.O.H. supervised the project.

## Supplementary information

**Supplementary Table 1.** Scripts used to process the “Narratives” data with brief descriptions. Scripts are listed roughly in order of execution. All scripts are located in the `code/` directory of the BIDS dataset and in the accompanying GitHub repository (https://github.com/snastase/narratives).

| Script | Description |
| --- | --- |
| `compile_metadata.py` | Compile dictionaries containing metadata and filenames for each subject (`subject_meta.json`) and task (`task_meta.json`). |
| `brain_masks.py` | Create brain masks for data in MNI space and fsaverage6 space. |
| `roi_masks.py` | Create early auditory cortex ROI masks based on multimodal cortical parcellation. |
| `run_pydeface.py` | Run PyDeface to anonymize (de-face) each subject’s anatomical image(s). |
| `run_mriqc.sh` | Run participant-level MRIQC on BIDS-formatted data for one subject. |
| `slurm_mriqc.sh` | Submit Slurm job array to run MRIQC on many subjects in parallel. |
| `run_mriqc_group.sh` | Run group-level MRIQC to summarize participant-level MRIQC outputs. |
| `run_fmriprep.sh` | Run fMRIPrep on BIDS-formatted data for one subject. |
| `slurm_fmriprep.sh` | Submit Slurm job array to run fMRIPrep on many subjects in parallel. |
| `run_smoothing.py` | Run spatial smoothing using AFNI's 3dBlurToFWHM and SurfSmooth for one subject. |
| `slurm_smoothing.sh` | Submit Slurm job array to run spatial smoothing on many subjects in parallel. |
| `extract_confounds.py` | Extract confound variables from fMRIPrep outputs for use with AFNI's 3dTproject. |
| `run_regression.py` | Run confound regression using AFNI's 3dTproject for one subject. |
| `slurm_regression.py` | Submit Slurm job array to run confound regression on many subjects in parallel. |
| `run_isc.py` | Run whole-brain vertex-wise leave-one-out intersubject correlation (ISC) analysis on smoothed surface data across all subjects in each story. |
| `roi_average.py` | Average non-smoothed time series across vertices within early auditory cortex ROI. |
| `roi_isc.py` | Compute leave-one-out ISC for early auditory cortex ROI. |
| `roi_lags.py` | Compute ISC for early auditory cortex ROI at lags ranging from -30 to +30 TRs. |
| `exclude_scans.py` | Compile dictionary (`scan_exclude.json`) for excluding scans based on ISC in early auditory cortex ROI with `exclude_scan` function. |
| `get_demog.py` | Summarize demographic information (age, sex) based on `participants.tsv` file |
| `get_words.py` | Summarize the number of words (including missing and unknown) across all stimuli. |
| `get_tsnr.py` | Estimate tSNR using AFNI's 3dTstat and compute median tSNR across all scans. |
| `get_fwhm.py` | Estimate intrinsic smoothness using AFNI's 3dFWHMx for one subject. |
| `slurm_fwhm.py` | Submit Slurm job array to run smoothness estimation on many subjects in parallel. |
| `plot_stim.py` | Plot audio waveform for Pie Man stimulus with example voxel time series (Figure 1). |
| `plot_qc.py` | Plot tSNR and FD from MRIQC, as well as intrinsic smoothness (Figure 2). |
| `plot_isc.py` | Plot ISC and lagged ISC for the early auditory cortex ROI (Figure 3). |
| `gifti_io.py` | Helper functions for reading and writing GIFTI surface files in Python. |

## References

1. Hasson, U., Ghazanfar, A. A., Galantucci, B., Garrod, S. & Keysers, C. Brain-to-brain coupling: a mechanism for creating and sharing a social world. Trends Cogn. Sci. 16, 114–121 (2012).

2. Berwick, R. C., Friederici, A. D., Chomsky, N. & Bolhuis, J. J. Evolution, brain, and the nature of language. Trends Cogn. Sci. 17, 89–98 (2013).

3. Bolhuis, J. J., Beckers, G. J. L., Huybregts, M. A. C., Berwick, R. C. & Everaert, M. B. H. Meaningful syntactic structure in songbird vocalizations? PLoS Biol. 16, e2005157 (2018).

4. Townsend, S. W., Engesser, S., Stoll, S., Zuberbühler, K. & Bickel, B. Compositionality in animals and humans. PLoS Biol. 16, e2006425 (2018).

5. Hamilton, L. S. & Huth, A. G. The revolution will not be controlled: natural stimuli in speech neuroscience. Lang. Cogn. Neurosci. 35, 573–582 (2020).

6. Hasson, U., Egidi, G., Marelli, M. & Willems, R. M. Grounding the neurobiology of language in first principles: The necessity of non-language-centric explanations for language comprehension. Cognition 180, 135–157 (2018).

7. Willems, R. M., Nastase, S. A. & Milivojevic, B. Narratives for neuroscience. Trends Neurosci. 43, 271–273 (2020).

8. Bookheimer, S. Functional MRI of language: new approaches to understanding the cortical organization of semantic processing. Annu. Rev. Neurosci. 25, 151–188 (2002).

9. Vigneau, M. et al. Meta-analyzing left hemisphere language areas: phonology, semantics, and sentence processing. Neuroimage 30, 1414–1432 (2006).

10. Hickok, G. & Poeppel, D. The cortical organization of speech processing. Nat. Rev. Neurosci. 8, 393–402 (2007).

11. Price, C. J. The anatomy of language: a review of 100 fMRI studies published in 2009. Ann. N. Y. Acad. Sci. 1191, 62–88 (2010).

12. Price, C. J. A review and synthesis of the first 20 years of PET and fMRI studies of heard speech, spoken language and reading. Neuroimage 62, 816–847 (2012).

13. Friederici, A. D. The brain basis of language processing: from structure to function. Physiol. Rev. 91, 1357–1392 (2011).

14. Friederici, A. D. The cortical language circuit: from auditory perception to sentence comprehension. Trends Cogn. Sci. 16, 262–268 (2012).

15. Kwong, K. K. et al. Dynamic magnetic resonance imaging of human brain activity during primary sensory stimulation. Proc. Natl. Acad. Sci. U. S. A. 89, 5675–5679 (1992).

16. Ogawa, S. et al. Intrinsic signal changes accompanying sensory stimulation: functional brain mapping with magnetic resonance imaging. Proc. Natl. Acad. Sci. U. S. A. 89, 5951–5955 (1992).

17. Logothetis, N. K., Pauls, J., Augath, M., Trinath, T. & Oeltermann, A. Neurophysiological investigation of the basis of the fMRI signal. Nature 412, 150–157 (2001).

18. Logothetis, N. K. What we can do and what we cannot do with fMRI. Nature 453, 869–878 (2008).

19. Démonet, J. F. et al. The anatomy of phonological and semantic processing in normal subjects. Brain 115, 1753–1768 (1992).

20. Zatorre, R. J., Evans, A. C., Meyer, E. & Gjedde, A. Lateralization of phonetic and pitch discrimination in speech processing. Science 256, 846–849 (1992).

21. Belin, P., Zatorre, R. J., Lafaille, P., Ahad, P. & Pike, B. Voice-selective areas in human auditory cortex. Nature 403, 309–312 (2000).

22. Vouloumanos, A., Kiehl, K. A., Werker, J. F. & Liddle, P. F. Detection of sounds in the auditory stream: event-related fMRI evidence for differential activation to speech and nonspeech. J. Cogn. Neurosci. 13, 994–1005 (2001).

23. Dapretto, M. & Bookheimer, S. Y. Form and content: dissociating syntax and semantics in sentence comprehension. Neuron 24, 427–432 (1999).

24. Ben-Shachar, M., Hendler, T., Kahn, I., Ben-Bashat, D. & Grodzinsky, Y. The neural reality of syntactic transformations: evidence from functional magnetic resonance imaging. Psychol. Sci. 14, 433–440 (2003).

25. Noppeney, U. & Price, C. J. An FMRI study of syntactic adaptation. J. Cogn. Neurosci. 16, 702–713 (2004).

26. Patterson, K., Nestor, P. J. & Rogers, T. T. Where do you know what you know? The representation of semantic knowledge in the human brain. Nat. Rev. Neurosci. 8, 976–987 (2007).

27. Binder, J. R., Desai, R. H., Graves, W. W. & Conant, L. L. Where is the semantic system? A critical review and meta-analysis of 120 functional neuroimaging studies. Cereb. Cortex 19, 2767–2796 (2009).

28. Fedorenko, E., Hsieh, P.-J., Nieto-Castañón, A., Whitfield-Gabrieli, S. & Kanwisher, N. New method for fMRI investigations of language: defining ROIs functionally in individual subjects. J. Neurophysiol. 104, 1177–1194 (2010).

29. Mahowald, K. & Fedorenko, E. Reliable individual-level neural markers of high-level language processing: a necessary precursor for relating neural variability to behavioral and genetic variability. Neuroimage 139, 74–93 (2016).

30. Braga, R. M., DiNicola, L. M., Becker, H. C. & Buckner, R. L. Situating the left-lateralized language network in the broader organization of multiple specialized large-scale distributed networks. J. Neurophysiol. 124, 1415–1448 (2020).

31. Jäncke, L., Wüstenberg, T., Scheich, H. & Heinze, H.-J. Phonetic perception and the temporal cortex. Neuroimage 15, 733–746 (2002).

32. Obleser, J., Zimmermann, J., Van Meter, J. & Rauschecker, J. P. Multiple stages of auditory speech perception reflected in event-related FMRI. Cereb. Cortex 17, 2251–2257 (2007).

33. Petersen, S. E., Fox, P. T., Posner, M. I., Mintun, M. & Raichle, M. E. Positron emission tomographic studies of the cortical anatomy of single-word processing. Nature 331, 585–589 (1988).

34. Wise, R. et al. Distribution of cortical neural networks involved in word comprehension and word retrieval. Brain 114, 1803–1817 (1991).

35. Poldrack, R. A. et al. Functional specialization for semantic and phonological processing in the left inferior prefrontal cortex. Neuroimage 10, 15–35 (1999).

36. Just, M. A., Carpenter, P. A., Keller, T. A., Eddy, W. F. & Thulborn, K. R. Brain activation modulated by sentence comprehension. Science 274, 114–116 (1996).

37. Kuperberg, G. R. et al. Common and distinct neural substrates for pragmatic, semantic, and syntactic processing of spoken sentences: an fMRI study. J. Cogn. Neurosci. 12, 321–341 (2000).

38. Ni, W. et al. An event-related neuroimaging study distinguishing form and content in sentence processing. J. Cogn. Neurosci. 12, 120–133 (2000).

39. Scott, S. K., Blank, C. C., Rosen, S. & Wise, R. J. Identification of a pathway for intelligible speech in the left temporal lobe. Brain 123, 2400–2406 (2000).

40. Vandenberghe, R., Nobre, A. C. & Price, C. J. The response of left temporal cortex to sentences. J. Cogn. Neurosci. 14, 550–560 (2002).

41. Humphries, C., Binder, J. R., Medler, D. A. & Liebenthal, E. Syntactic and semantic modulation of neural activity during auditory sentence comprehension. J. Cogn. Neurosci. 18, 665–679 (2006).

42. Brennan, J. et al. Syntactic structure building in the anterior temporal lobe during natural story listening. Brain Lang. 120, 163–173 (2012).

43. Brennan, J. R., Stabler, E. P., Van Wagenen, S. E., Luh, W.-M. & Hale, J. T. Abstract linguistic structure correlates with temporal activity during naturalistic comprehension. Brain Lang. 157-158, 81–94 (2016).

44. Nastase, S. A., Goldstein, A. & Hasson, U. Keep it real: rethinking the primacy of experimental control in cognitive neuroscience. Neuroimage 222, 117254 (2020).

45. Wehbe, L. et al. Simultaneously uncovering the patterns of brain regions involved in different story reading subprocesses. PLoS One 9, e112575 (2014).

46. Huth, A. G., de Heer, W. A., Griffiths, T. L., Theunissen, F. E. & Gallant, J. L. Natural speech reveals the semantic maps that tile human cerebral cortex. Nature 532, 453–458 (2016).

47. Goldberg, Y. Neural network methods for natural language processing. Synth. Lectures Hum. Lang. Technol. 10, 1–309 (2017).

48. Baroni, M. Linguistic generalization and compositionality in modern artificial neural networks. Philos. Trans. R. Soc. Lond. B Biol. Sci. 375, 20190307 (2020).

49. Devlin, J., Chang, M.-W., Lee, K. & Toutanova, K. BERT: pre-training of deep bidirectional transformers for language understanding. arXiv (2018).

50. Radford, A. et al. Language models are unsupervised multitask learners. OpenAI Blog (2019).

51. Turney, P. D. & Pantel, P. From frequency to meaning: vector space models of semantics. J. Artif. Intell. Res. 37, 141–188 (2010).

52. Mikolov, T., Sutskever, I., Chen, K., Corrado, G. S. & Dean, J. Distributed representations of words and phrases and their compositionality. in Advances in Neural Information Processing Systems 26 (eds. Burges, C. J. C., Bottou, L., Welling, M., Ghahramani, Z. & Weinberger, K. Q.) 3111–3119 (Curran Associates, Inc., 2013).

53. Manning, C. D., Clark, K., Hewitt, J., Khandelwal, U. & Levy, O. Emergent linguistic structure in artificial neural networks trained by self-supervision. Proc. Natl. Acad. Sci. U. S. A. 117, 30046–30054 (2020).

54. Breiman, L. Statistical modeling: the two cultures. Stat. Sci. 16, 199–231 (2001).

55. Yarkoni, T. & Westfall, J. Choosing prediction over explanation in psychology: lessons from machine learning. Perspect. Psychol. Sci. 12, 1100–1122 (2017).

56. Varoquaux, G. & Poldrack, R. A. Predictive models avoid excessive reductionism in cognitive neuroimaging. Curr. Opin. Neurobiol. 55, 1–6 (2019).

57. Hasson, U., Nastase, S. A. & Goldstein, A. Direct fit to nature: an evolutionary perspective on biological and artificial neural networks. Neuron 105, 416–434 (2020).

58. LeCun, Y., Cortes, C. & Burges, C. J. MNIST handwritten digit database. (2010).

59. Krizhevsky, A. Learning multiple layers of features from tiny images. (University of Toronto, 2009).

60. Milham, M. P. et al. Assessment of the impact of shared brain imaging data on the scientific literature. Nat. Commun. 9, 2818 (2018).

61. Biswal, B. B. et al. Toward discovery science of human brain function. Proc. Natl. Acad. Sci. U. S. A. 107, 4734–4739 (2010).

62. Van Essen, D. C. et al. The WU-Minn Human Connectome Project: an overview. Neuroimage 80, 62–79 (2013).

63. Shafto, M. A. et al. The Cambridge Centre for Ageing and Neuroscience (Cam-CAN) study protocol: a cross-sectional, lifespan, multidisciplinary examination of healthy cognitive ageing. BMC Neurol. 14, 204 (2014).

64. Taylor, J. R. et al. The Cambridge Centre for Ageing and Neuroscience (Cam-CAN) data repository: structural and functional MRI, MEG, and cognitive data from a cross-sectional adult lifespan sample. Neuroimage 144, 262–269 (2017).

65. Alexander, L. M. et al. An open resource for transdiagnostic research in pediatric mental health and learning disorders. Sci Data 4, 170181 (2017).

66. Poldrack, R. A. & Gorgolewski, K. J. Making big data open: data sharing in neuroimaging. Nat. Neurosci. 17, 1510–1517 (2014).

67. Poldrack, R. A. et al. Scanning the horizon: towards transparent and reproducible neuroimaging research. Nat. Rev. Neurosci. 18, 115–126 (2017).

68. Poldrack, R. A., Gorgolewski, K. J. & Varoquaux, G. Computational and informatic advances for reproducible data analysis in neuroimaging. Annu. Rev. Biomed. Data Sci. 2, 119–138 (2019).

69. Ferguson, A. R., Nielson, J. L., Cragin, M. H., Bandrowski, A. E. & Martone, M. E. Big data from small data: data-sharing in the ‘long tail’ of neuroscience. Nat. Neurosci. 17, 1442–1447 (2014).

70. Hanke, M. et al. A high-resolution 7-Tesla fMRI dataset from complex natural stimulation with an audio movie. Sci Data 1, 140003 (2014).

71. Hanke, M. et al. A studyforrest extension, simultaneous fMRI and eye gaze recordings during prolonged natural stimulation. Sci Data 3, 160092 (2016).

72. Aly, M., Chen, J., Turk-Browne, N. B. & Hasson, U. Learning naturalistic temporal structure in the posterior medial network. J. Cogn. Neurosci. 30, 1345–1365 (2018).

73. DuPre, E., Hanke, M. & Poline, J.-B. Nature abhors a paywall: how open science can realize the potential of naturalistic stimuli. Neuroimage 216, 116330 (2020).

74. Aliko, S., Huang, J., Gheorghiu, F., Meliss, S. & Skipper, J. I. A naturalistic neuroimaging database for understanding the brain using ecological stimuli. Sci Data 7, 347 (2020).

75. Richardson, H., Lisandrelli, G., Riobueno-Naylor, A. & Saxe, R. Development of the social brain from age three to twelve years. Nat. Commun. 9, 1027 (2018).

76. Finn, E. S., Corlett, P. R., Chen, G., Bandettini, P. A. & Constable, R. T. Trait paranoia shapes inter-subject synchrony in brain activity during an ambiguous social narrative. Nat. Commun. 9, 2043 (2018).

77. Chen, J. et al. Accessing real-life episodic information from minutes versus hours earlier modulates hippocampal and high-order cortical dynamics. Cereb. Cortex 26, 3428–3441 (2016).

78. Chen, J. et al. Shared memories reveal shared structure in neural activity across individuals. Nat. Neurosci. 20, 115–125 (2017).

79. O’Connor, D. et al. The Healthy Brain Network Serial Scanning Initiative: a resource for evaluating inter-individual differences and their reliabilities across scan conditions and sessions. GigaScience 6, 1–14 (2017).

80. Haxby, J. V. et al. A common, high-dimensional model of the representational space in human ventral temporal cortex. Neuron 72, 404–416 (2011).

81. Nastase, S. A. et al. Attention Selectively Reshapes the Geometry of Distributed Semantic Representation. Cereb. Cortex 27, 4277–4291 (2017).

82. Nastase, S. A., Halchenko, Y. O., Connolly, A. C., Gobbini, M. I. & Haxby, J. V. Neural responses to naturalistic clips of behaving animals in two different task contexts. Front. Neurosci. 12, 316 (2018).

83. Castello, M. V. di O., di Oleggio Castello, M. V., Chauhan, V., Jiahui, G. & Ida Gobbini, M. An fMRI dataset in response to ‘The Grand Budapest Hotel’, a socially-rich, naturalistic movie. Scientific Data vol. 7 (2020).

84. Nastase, S. A. et al. Narratives. (2019) doi:10.18112/OPENNEURO.DS002345.V1.0.1.

85. Gorgolewski, K. J. et al. The brain imaging data structure, a format for organizing and describing outputs of neuroimaging experiments. Sci Data 3, 160044 (2016).

86. Poldrack, R. A. & Gorgolewski, K. J. OpenfMRI: Open sharing of task fMRI data. Neuroimage 144, 259–261 (2017).

87. Hanke, M. et al. datalad/datalad: 0.13.3 (August 28, 2020). (2020). doi:10.5281/zenodo.4006562.

88. Hanke, M. et al. In defense of decentralized research data management. Neuroforum 27, 17–25 (2021).

89. Spiers, H. J. & Maguire, E. A. Decoding human brain activity during real-world experiences. Trends Cogn. Sci. 11, 356–365 (2007).

90. Hasson, U. & Honey, C. J. Future trends in neuroimaging: neural processes as expressed within real-life contexts. Neuroimage 62, 1272–1278 (2012).

91. Matusz, P. J., Dikker, S., Huth, A. G. & Perrodin, C. Are we ready for real-world neuroscience? J. Cogn. Neurosci. 31, 327–338 (2019).

92. Sonkusare, S., Breakspear, M. & Guo, C. Naturalistic stimuli in neuroscience: critically acclaimed. Trends Cogn. Sci. 23, 699–714 (2019).

93. Redcay, E. & Moraczewski, D. Social cognition in context: a naturalistic imaging approach. Neuroimage 216, 116392 (2020).

94. Vanderwal, T., Eilbott, J. & Castellanos, F. X. Movies in the magnet: naturalistic paradigms in developmental functional neuroimaging. Dev. Cogn. Neurosci. 36, 100600 (2018).

95. Kriegeskorte, N., Mur, M. & Bandettini, P. A. Representational similarity analysis–connecting the branches of systems neuroscience. Front. Syst. Neurosci. 2, 4 (2008).

96. Naselaris, T., Kay, K. N., Nishimoto, S. & Gallant, J. L. Encoding and decoding in fMRI. Neuroimage 56, 400–410 (2011).

97. Santoro, R. et al. Encoding of natural sounds at multiple spectral and temporal resolutions in the human auditory cortex. PLoS Comput. Biol. 10, e1003412 (2014).

98. de Heer, W. A., Huth, A. G., Griffiths, T. L., Gallant, J. L. & Theunissen, F. E. The hierarchical cortical organization of human speech processing. J. Neurosci. 37, 6539–6557 (2017).

99. Kell, A. J. E., Yamins, D. L. K., Shook, E. N., Norman-Haignere, S. V. & McDermott, J. H. A task-optimized neural network replicates human auditory behavior, predicts brain responses, and reveals a cortical processing hierarchy. Neuron 98, 630–644.e16 (2018).

100. Mitchell, T. M. et al. Predicting human brain activity associated with the meanings of nouns. Science 320, 1191–1195 (2008).

101. Pereira, F. et al. Toward a universal decoder of linguistic meaning from brain activation. Nat. Commun. 9, 963 (2018).

102. Schrimpf, M. et al. The neural architecture of language: integrative reverse-engineering converges on a model for predictive processing. bioRxiv (2020) doi:10.1101/2020.06.26.174482.

103. Hasson, U., Nir, Y., Levy, I., Fuhrmann, G. & Malach, R. Intersubject synchronization of cortical activity during natural vision. Science 303, 1634–1640 (2004).

104. Nastase, S. A., Gazzola, V., Hasson, U. & Keysers, C. Measuring shared responses across subjects using intersubject correlation. Soc. Cogn. Affect. Neurosci. 14, 667–685 (2019).

105. Vanderwal, T. et al. Individual differences in functional connectivity during naturalistic viewing conditions. Neuroimage 157, 521–530 (2017).

106. Feilong, M., Nastase, S. A., Guntupalli, J. S. & Haxby, J. V. Reliable individual differences in fine-grained cortical functional architecture. Neuroimage 183, 375–386 (2018).

107. Finn, E. S. et al. Idiosynchrony: from shared responses to individual differences during naturalistic neuroimaging. Neuroimage 215, 116828 (2020).

108. Chen, P.-H. et al. A reduced-dimension fMRI shared response model. in Advances in Neural Information Processing Systems 28 (eds. Cortes, C., Lawrence, N. D., Lee, D. D., Sugiyama, M. & Garnett, R.) 460–468 (Curran Associates, Inc., 2015).

109. Guntupalli, J. S. et al. A model of representational spaces in human cortex. Cereb. Cortex 26, 2919–2934 (2016).

110. Guntupalli, J. S., Feilong, M. & Haxby, J. V. A computational model of shared fine-scale structure in the human connectome. PLoS Comput. Biol. 14, e1006120 (2018).

111. Van Uden, C. E. et al. Modeling semantic encoding in a common neural representational space. Front. Neurosci. 12, 437 (2018).

112. Haxby, J. V., Guntupalli, J. S., Nastase, S. A. & Feilong, M. Hyperalignment: modeling shared information encoded in idiosyncratic cortical topographies. eLife 9, (2020).

113. Milivojevic, B., Varadinov, M., Vicente Grabovetsky, A., Collin, S. H. P. & Doeller, C. F. Coding of event nodes and narrative context in the hippocampus. J. Neurosci. 36, 12412–12424 (2016).

114. Baldassano, C. et al. Discovering event structure in continuous narrative perception and memory. Neuron 95, 709–721.e5 (2017).

115. Baldassano, C., Hasson, U. & Norman, K. A. Representation of real-world event schemas during narrative perception. J. Neurosci. 38, 9689–9699 (2018).

116. Chang, L. J. et al. Endogenous variation in ventromedial prefrontal cortex state dynamics during naturalistic viewing reflects affective experience. Sci. Adv. 7, eabf7129 (2021).

117. Heusser, A. C., Fitzpatrick, P. C. & Manning, J. R. Geometric models reveal behavioural and neural signatures of transforming experiences into memories. Nat Hum Behav (2021) doi:10.1038/s41562-021-01051-6.

118. Simony, E. et al. Dynamic reconfiguration of the default mode network during narrative comprehension. Nat. Commun. 7, 12141 (2016).

119. Kim, D., Kay, K., Shulman, G. L. & Corbetta, M. A new modular brain organization of the BOLD signal during natural vision. Cereb. Cortex 28, 3065–3081 (2018).

120. Betzel, R. F., Byrge, L., Esfahlani, F. Z. & Kennedy, D. P. Temporal fluctuations in the brain’s modular architecture during movie-watching. Neuroimage 213, 116687 (2020).

121. Meer, J. N. van der, Breakspear, M., Chang, L. J., Sonkusare, S. & Cocchi, L. Movie viewing elicits rich and reliable brain state dynamics. Nat. Commun. 11, 5004 (2020).

122. Brainard, D. H. The Psychophysics Toolbox. Spat. Vis. 10, 433–436 (1997).

123. Kleiner, M., Brainard, D. & Pelli, D. What’s new in Psychtoolbox-3? Perception 36 ECVP Abstract Supplement (2007).

124. Peirce, J. W. PsychoPy—psychophysics software in Python. J. Neurosci. Methods 162, 8–13 (2007).

125. Peirce, J. W. Generating stimuli for neuroscience using PsychoPy. Front. Neuroinform. 2, 10 (2009).

126. Peirce, J. et al. PsychoPy2: experiments in behavior made easy. Behav. Res. Methods 51, 195–203 (2019).

127. DuPre, E., Hanke, M. & Poline, J.-B. Nature abhors a paywall: how open science can realize the potential of naturalistic stimuli. (2019) doi:10.31234/osf.io/sdbqv.

128. McNamara, Q., De La Vega, A. & Yarkoni, T. Developing a comprehensive framework for multimodal feature extraction. in Proceedings of the 23rd ACM SIGKDD International Conference on Knowledge Discovery and Data Mining 1567–1574 (ACM, 2017). doi:10.1145/3097983.3098075.

129. Ochshorn, R. M. & Hawkins, M. Gentle: a robust yet lenient forced aligner built on Kaldi. (2016).

130. Povey, D. et al. The Kaldi speech recognition toolkit. in IEEE 2011 workshop on automatic speech recognition and understanding (IEEE Signal Processing Society, 2011).

131. Cieri, C., Miller, D. & Walker, K. The Fisher Corpus: a resource for the next generations of speech-to-text. in Proceedings of the Fourth International Conference on Language Resources and Evaluation (LREC) vol. 4 69–71 (2004).

132. Nichols, T. E. et al. Best practices in data analysis and sharing in neuroimaging using MRI. Nat. Neurosci. 20, 299–303 (2017).

133. Gulban, O. F. et al. poldracklab/pydeface: v2.0.0. (2019). doi:10.5281/zenodo.3524401.

134. Esteban, O. et al. fMRIPrep: a robust preprocessing pipeline for functional MRI. Nat. Methods 16, 111–116 (2019).

135. Esteban, O. et al. fMRIPrep: a robust preprocessing pipeline for functional MRI. (2020). doi:10.5281/zenodo.3724468.

136. Gorgolewski, K. et al. Nipype: a flexible, lightweight and extensible neuroimaging data processing framework in python. Front. Neuroinform. 5, 13 (2011).

137. Esteban, O. et al. nipy/nipype: 1.4.2. (2020). doi:10.5281/zenodo.3668316.

138. Abraham, A. et al. Machine learning for neuroimaging with scikit-learn. Front. Neuroinform. 8, 14 (2014).

139. Kurtzer, G. M., Sochat, V. & Bauer, M. W. Singularity: scientific containers for mobility of compute. PLoS One 12, e0177459 (2017).

140. Cox, R. W. AFNI: software for analysis and visualization of functional magnetic resonance neuroimages. Comput. Biomed. Res. 29, 162–173 (1996).

141. Cox, R. W. AFNI: what a long strange trip it’s been. Neuroimage 62, 743–747 (2012).

142. Tustison, N. J. et al. N4ITK: improved N3 bias correction. IEEE Trans. Med. Imaging 29, 1310–1320 (2010).

143. Avants, B. B., Epstein, C. L., Grossman, M. & Gee, J. C. Symmetric diffeomorphic image registration with cross-correlation: evaluating automated labeling of elderly and neurodegenerative brain. Med. Image Anal. 12, 26–41 (2008).

144. Zhang, Y., Brady, M. & Smith, S. Segmentation of brain MR images through a hidden Markov random field model and the expectation-maximization algorithm. IEEE Trans. Med. Imaging 20, 45–57 (2001).

145. Dale, A. M., Fischl, B. & Sereno, M. I. Cortical surface-based analysis. I. Segmentation and surface reconstruction. Neuroimage 9, 179–194 (1999).

146. Fischl, B. FreeSurfer. Neuroimage 62, 774–781 (2012).

147. Klein, A. et al. Mindboggling morphometry of human brains. PLoS Comput. Biol. 13, e1005350 (2017).

148. Esteban, O., Ciric, R., Markiewicz, C. J., Poldrack, R. A. & Gorgolewski, K. J. TemplateFlow Client: accessing the library of standardized neuroimaging standard spaces. (2020). doi:10.5281/zenodo.3981009.

149. Fonov, V. S., Evans, A. C., McKinstry, R. C., Almli, C. R. & Collins, D. L. Unbiased nonlinear average age-appropriate brain templates from birth to adulthood. Neuroimage 47, S102 (2009).

150. Evans, A. C., Janke, A. L., Collins, D. L. & Baillet, S. Brain templates and atlases. Neuroimage 62, 911–922 (2012).

151. Fischl, B., Sereno, M. I., Tootell, R. B. & Dale, A. M. High-resolution intersubject averaging and a coordinate system for the cortical surface. Hum. Brain Mapp. 8, 272–284 (1999).

152. Huntenburg, J. M. Evaluating nonlinear coregistration of BOLD EPI and T1w images. (Freie Universität Berlin, 2014).

153. Wang, S. et al. Evaluation of Field Map and Nonlinear Registration Methods for Correction of Susceptibility Artifacts in Diffusion MRI. Front. Neuroinform. 11, 17 (2017).

154. Treiber, J. M. et al. Characterization and Correction of Geometric Distortions in 814 Diffusion Weighted Images. PLoS One 11, e0152472 (2016).

155. Greve, D. N. & Fischl, B. Accurate and robust brain image alignment using boundary-based registration. Neuroimage 48, 63–72 (2009).

156. Jenkinson, M., Bannister, P., Brady, M. & Smith, S. Improved optimization for the robust and accurate linear registration and motion correction of brain images. Neuroimage 17, 825–841 (2002).

157. Smith, S. M. et al. Advances in functional and structural MR image analysis and implementation as FSL. Neuroimage 23, S208–19 (2004).

158. Jenkinson, M., Beckmann, C. F., Behrens, T. E. J., Woolrich, M. W. & Smith, S. M. FSL. Neuroimage 62, 782–790 (2012).

159. Cox, R. W. & Hyde, J. S. Software tools for analysis and visualization of fMRI data. NMR Biomed. 10, 171–178 (1997).

160. Lanczos, C. Evaluation of Noisy Data. J. Soc. Ind. Appl. Math. B Numer. Anal. 1, 76–85 (1964).

161. Power, J. D. et al. Methods to detect, characterize, and remove motion artifact in resting state fMRI. Neuroimage 84, 320–341 (2014).

162. Behzadi, Y., Restom, K., Liau, J. & Liu, T. T. A component based noise correction method (CompCor) for BOLD and perfusion based fMRI. Neuroimage 37, 90–101 (2007).

163. Satterthwaite, T. D. et al. An improved framework for confound regression and filtering for control of motion artifact in the preprocessing of resting-state functional connectivity data. Neuroimage 64, 240–256 (2013).

164. Pajula, J. & Tohka, J. Effects of spatial smoothing on inter-subject correlation based analysis of FMRI. Magn. Reson. Imaging 32, 1114–1124 (2014).

165. Nastase, S. A., Liu, Y.-F., Hillman, H., Norman, K. A. & Hasson, U. Leveraging shared connectivity to aggregate heterogeneous datasets into a common response space. Neuroimage 217, 116865 (2020).

166. Chung, M. K. et al. Cortical thickness analysis in autism with heat kernel smoothing. Neuroimage 25, 1256–1265 (2005).

167. Hagler, D. J., Jr, Saygin, A. P. & Sereno, M. I. Smoothing and cluster thresholding for cortical surface-based group analysis of fMRI data. Neuroimage 33, 1093–1103 (2006).

168. Triantafyllou, C., Hoge, R. D. & Wald, L. L. Effect of spatial smoothing on physiological noise in high-resolution fMRI. Neuroimage 32, 551–557 (2006).

169. Friedman, L., Glover, G. H., Krenz, D., Magnotta, V. & FIRST BIRN. Reducing inter-scanner variability of activation in a multicenter fMRI study: role of smoothness equalization. Neuroimage 32, 1656–1668 (2006).

170. Simony, E. & Chang, C. Analysis of stimulus-induced brain dynamics during naturalistic paradigms. Neuroimage 216, 116461 (2019).

171. Ciric, R. et al. Benchmarking of participant-level confound regression strategies for the control of motion artifact in studies of functional connectivity. Neuroimage 154, 174–187 (2017).

172. Parkes, L., Fulcher, B., Yücel, M. & Fornito, A. An evaluation of the efficacy, reliability, and sensitivity of motion correction strategies for resting-state functional MRI. Neuroimage 171, 415–436 (2018).

173. Muschelli, J. et al. Reduction of motion-related artifacts in resting state fMRI using aCompCor. Neuroimage 96, 22–35 (2014).

174. Lindquist, M. A., Geuter, S., Wager, T. D. & Caffo, B. S. Modular preprocessing pipelines can reintroduce artifacts into fMRI data. Hum. Brain Mapp. 40, 2358–2376 (2019).

175. Halchenko, Y. O. & Hanke, M. Open is not enough. Let’s take the next step: an integrated, community-driven computing platform for neuroscience. Front. Neuroinform. 6, 22 (2012).

176. Hanke, M. & Halchenko, Y. O. Neuroscience runs on GNU/Linux. Front. Neuroinform. 5, 8 (2011).

177. Walt, S. van der, Colbert, S. C. & Varoquaux, G. The NumPy Array: a structure for efficient numerical computation. Comput. Sci. Eng. 13, 22–30 (2011).

178. Harris, C. R. et al. Array programming with NumPy. Nature 585, 357–362 (2020).

179. Jones, E., Oliphant, T. & Peterson, P. SciPy: open source scientific tools for Python. (2001--).

180. Virtanen, P. et al. SciPy 1.0: fundamental algorithms for scientific computing in Python. Nat. Methods 17, 261–272 (2020).

181. McKinney, W. Data structures for statistical computing in Python. in Proceedings of the 9th Python in Science Conference 51–56 (2010).

182. Brett, M. et al. nipy/nibabel: 3.1.1. (2020). doi:10.5281/zenodo.3924343.

183. Perez, F. & Granger, B. E. IPython: a system for interactive scientific computing. Computing in Science Engineering 9, 21–29 (2007).

184. Kluyver, T. et al. Jupyter Notebooks—a publishing format for reproducible computational workflows. in Positioning and Power in Academic Publishing: Players, Agents and Agendas (eds. Loizides, F. & Schmidt, B.) 87–90 (IOS Press, 2016).

185. Jette, M. A., Yoo, A. B. & Grondona, M. SLURM: Simple Linux Utility for Resource Management. in Job Scheduling Strategies for Parallel Processing (eds. Feitelson, D., Rudolph, L. & Schwiegelshohn, U.) 44–60 (Springer, Berlin, Heidelberg, 2003).

186. Saad, Z. S., Reynolds, R. C., Argall, B., Japee, S. & Cox, R. W. SUMA: an interface for surface-based intra- and inter-subject analysis with AFNI. in 2004 2nd IEEE International Symposium on Biomedical Imaging: Nano to Macro (IEEE Cat No. 04EX821) 1510–1513 Vol. 2 (ieeexplore.ieee.org, 2004). doi:10.1109/ISBI.2004.1398837.

187. Saad, Z. S. & Reynolds, R. C. SUMA. Neuroimage 62, 768–773 (2012).

188. Hunter, J. D. Matplotlib: A 2D Graphics Environment. Comput. Sci. Eng. 9, 90–95 (2007).

189. Lerner, Y., Honey, C. J., Silbert, L. J. & Hasson, U. Topographic mapping of a hierarchy of temporal receptive windows using a narrated story. J. Neurosci. 31, 2906–2915 (2011).

190. Ben-Yakov, A., Honey, C. J., Lerner, Y. & Hasson, U. Loss of reliable temporal structure in event-related averaging of naturalistic stimuli. Neuroimage 63, 501–506 (2012).

191. Regev, M., Honey, C. J., Simony, E. & Hasson, U. Selective and invariant neural responses to spoken and written narratives. J. Neurosci. 33, 15978–15988 (2013).

192. Stephens, G. J., Honey, C. J. & Hasson, U. A place for time: the spatiotemporal structure of neural dynamics during natural audition. J. Neurophysiol. 110, 2019–2026 (2013).

193. Lerner, Y., Honey, C. J., Katkov, M. & Hasson, U. Temporal scaling of neural responses to compressed and dilated natural speech. J. Neurophysiol. 111, 2433–2444 (2014).

194. Liu, Y. et al. Measuring speaker-listener neural coupling with functional near infrared spectroscopy. Sci. Rep. 7, 43293 (2017).

195. Vodrahalli, K. et al. Mapping between fMRI responses to movies and their natural language annotations. Neuroimage 180, 223–231 (2018).

196. Yeshurun, Y., Nguyen, M. & Hasson, U. Amplification of local changes along the timescale processing hierarchy. Proc. Natl. Acad. Sci. U. S. A. 114, 9475–9480 (2017).

197. Zuo, X. et al. Temporal integration of narrative information in a hippocampal amnesic patient. Neuroimage 213, 116658 (2020).

198. Gross, J. et al. Speech rhythms and multiplexed oscillatory sensory coding in the human brain. PLoS Biol. 11, e1001752 (2013).

199. Blank, I. A. & Fedorenko, E. Domain-general brain regions do not track linguistic input as closely as language-selective regions. J. Neurosci. 37, 9999–10011 (2017).

200. Iotzov, I. et al. Divergent neural responses to narrative speech in disorders of consciousness. Ann Clin Transl Neurol 4, 784–792 (2017).

201. Loiotile, R. E., Cusack, R. & Bedny, M. Naturalistic audio-movies and narrative synchronize ‘visual’ cortices across congenitally blind but not sighted individuals. J. Neurosci. 39, 8940–8948 (2019).

202. Lositsky, O. et al. Neural pattern change during encoding of a narrative predicts retrospective duration estimates. Elife 5, (2016).

203. Yeshurun, Y. et al. Same story, different story: the neural representation of interpretive frameworks. Psychol. Sci. 28, 307–319 (2017).

204. Regev, M. et al. Propagation of Information Along the Cortical Hierarchy as a Function of Attention While Reading and Listening to Stories. Cereb. Cortex 29, 4017–4034 (2019).

205. Chien, H.-Y. S. & Honey, C. J. Constructing and forgetting temporal context in the human cerebral cortex. Neuron 106, 675–686.e11 (2020).

206. Zadbood, A., Chen, J., Leong, Y. C., Norman, K. A. & Hasson, U. How we transmit memories to other brains: constructing shared neural representations via communication. Cereb. Cortex 27, 4988–5000 (2017).

207. Heider, F. & Simmel, M. An experimental study of apparent behavior. Am. J. Psychol. 57, 243–259 (1944).

208. Nguyen, M., Vanderwal, T. & Hasson, U. Shared understanding of narratives is correlated with shared neural responses. Neuroimage 184, 161–170 (2019).

209. Chang, C. H. C., Lazaridi, C., Yeshurun, Y., Norman, K. A. & Hasson, U. Relating the past with the present: Information integration and segregation during ongoing narrative processing. J. Cogn. Neurosci. 33, 1106–1128 (2021).

210. Visconti di Oleggio Castello, M. et al. ReproNim/reproin 0.6.0. (2020). doi:10.5281/zenodo.3625000.

211. Halchenko, Y. et al. nipy/heudiconv v0.8.0. (2020). doi:10.5281/zenodo.3760062.

212. Lin, X. et al. Data-efficient mutual information neural estimator. arXiv (2019).

213. Mennes, M., Biswal, B. B., Castellanos, F. X. & Milham, M. P. Making data sharing work: the FCP/INDI experience. Neuroimage 82, 683–691 (2013).

214. Kennedy, D. N., Haselgrove, C., Riehl, J., Preuss, N. & Buccigrossi, R. The NITRC image repository. Neuroimage 124, 1069–1073 (2016).

215. Wilkinson, M. D. et al. The FAIR Guiding Principles for scientific data management and stewardship. Sci Data 3, 160018 (2016).

216. Gorgolewski, K. J. et al. BIDS apps: improving ease of use, accessibility, and reproducibility of neuroimaging data analysis methods. PLoS Comput. Biol. 13, e1005209 (2017).

217. Cox, R. W. et al. A (sort of) new image data format standard: NIfTI-1. in 10th Annual Meeting of the Organization for Human Brain Mapping, Budapest, Hungary (2004).

218. Li, X., Morgan, P. S., Ashburner, J., Smith, J. & Rorden, C. The first step for neuroimaging data analysis: DICOM to NIfTI conversion. J. Neurosci. Methods 264, 47–56 (2016).

219. Wagner, A. S. et al. The DataLad Handbook. (2020). doi:10.5281/zenodo.3905791.

220. Esteban, O. et al. MRIQC: advancing the automatic prediction of image quality in MRI from unseen sites. PLoS One 12, e0184661 (2017).

221. Esteban, O. et al. MRIQC: advancing the automatic prediction of image quality in MRI from unseen sites. (2019). doi:10.5281/zenodo.3352432.

222. Power, J. D., Barnes, K. A., Snyder, A. Z., Schlaggar, B. L. & Petersen, S. E. Spurious but systematic correlations in functional connectivity MRI networks arise from subject motion. Neuroimage 59, 2142–2154 (2012).

223. Forman, S. D. et al. Improved assessment of significant activation in functional magnetic resonance imaging (fMRI): use of a cluster-size threshold. Magn. Reson. Med. 33, 636–647 (1995).

224. Krüger, G. & Glover, G. H. Physiological noise in oxygenation-sensitive magnetic resonance imaging. Magn. Reson. Med. 46, 631–637 (2001).

225. Ojemann, J. G. et al. Anatomic localization and quantitative analysis of gradient refocused echo-planar fMRI susceptibility artifacts. Neuroimage 6, 156–167 (1997).

226. Hasson, U., Malach, R. & Heeger, D. J. Reliability of cortical activity during natural stimulation. Trends Cogn. Sci. 14, 40–48 (2010).

227. Nili, H. et al. A toolbox for representational similarity analysis. PLoS Comput. Biol. 10, e1003553 (2014).

228. Silver, N. C. & Dunlap, W. P. Averaging correlation coefficients: should Fisher’s z transformation be used? Journal of Applied Psychology 72, 146–148 (1987).

229. Cohen, J. D. et al. Computational approaches to fMRI analysis. Nat. Neurosci. 20, 304–313 (2017).

230. Kumar, M. et al. BrainIAK tutorials: user-friendly learning materials for advanced fMRI analysis. PLoS Comp. Biol. (2020) doi:10.31219/osf.io/j4sbc.

231. Kumar, M. et al. BrainIAK: The Brain Imaging Analysis Kit. OSF Preprints (2020) doi:10.31219/osf.io/db2ev.

232. Glasser, M. F. et al. A multi-modal parcellation of human cerebral cortex. Nature 536, 171–178 (2016).

233. Mills, K. HCP-MMP1.0 projected on fsaverage. (2016) doi:10.6084/M9.FIGSHARE.3498446.V2.

234. Aguirre, G. K., Zarahn, E. & D’esposito, M. The variability of human, BOLD hemodynamic responses. Neuroimage 8, 360–369 (1998).

235. Handwerker, D. A., Ollinger, J. M. & D’Esposito, M. Variation of BOLD hemodynamic responses across subjects and brain regions and their effects on statistical analyses. Neuroimage 21, 1639–1651 (2004).

236. Binder, J. R. et al. Human temporal lobe activation by speech and nonspeech sounds. Cereb. Cortex 10, 512–528 (2000).

237. Zatorre, R. J., Belin, P. & Penhune, V. B. Structure and function of auditory cortex: music and speech. Trends Cogn. Sci. 6, 37–46 (2002).

238. Chen, G. et al. Untangling the relatedness among correlations, part I: nonparametric approaches to inter-subject correlation analysis at the group level. Neuroimage 142, 248–259 (2016).

239. Chen, G., Taylor, P. A., Shin, Y.-W., Reynolds, R. C. & Cox, R. W. Untangling the relatedness among correlations, Part II: inter-subject correlation group analysis through linear mixed-effects modeling. Neuroimage 147, 825–840 (2017).

240. Chen, G. et al. Untangling the relatedness among correlations, part III: inter-subject correlation analysis through Bayesian multilevel modeling for naturalistic scanning. Neuroimage 216, 116474 (2020).

241. Markiewicz, C. J. et al. poldracklab/fitlins. (Zenodo, 2021). doi:10.5281/zenodo.1306215.

242. de la Vega, A., Blair, R. & Yarkoni, T. neuroscout/neuroscout. (Zenodo, 2021). doi:10.5281/zenodo.2592674.

243. Yarkoni, T. et al. PyBIDS: Python tools for BIDS datasets. J. Open Source Softw. 4, (2019).

